# Systematic analysis of ADP-ribose detection reagents and optimisation of sample preparation to detect ADP-ribosylation *in vitro* and in cells

**DOI:** 10.1101/2022.02.22.481411

**Authors:** Lisa Weixler, Jim Voorneveld, Gülcan Aydin, Timo M. H. R. Bolte, Jeffrey Momoh, Mareike Bütepage, Alexandra Golzmann, Bernhard Lüscher, Dmitri V. Filippov, Roko Žaja, Karla L. H. Feijs

## Abstract

Recent evidence suggests that modification of substrates with a single ADP-ribose (ADPr) is important in for example antiviral immunity and cancer. However, the endogenous substrates and the extent of mono-ADP-ribosylation are still largely unclear. Several reagents were developed to detect ADP-ribosylation but it is unknown whether they recognise only ADPr, amino acid-ADPr linkages or a combination of ADPr with a protein backbone. We screened the affinity of selected reagents for enzymatically, chemically and in cell generated ADP-ribosylation on glutamate, cysteine, serine, arginine, threonine and RNA by blotting, as well as analysed the subcellular sites of ADP-ribosylation using immunofluorescence confocal microscopy. We furthermore observed that the modification is heat-labile and optimised sample preparation procedures. Our comparison of the available reagents, as well as optimisation of sample preparation, will allow future work further dissecting the function of ADP-ribosylation in cells, both on protein and on RNA substrates.

## INTRODUCTION

The posttranslational modification of proteins is a well-known way to regulate proteins in response to changes in nutrient availability, viral infection, DNA damage and many other signals. Proteins with catalytic activity can be switched on or off, others can change their localisation within the cell or interact with different molecules. ADP-ribosylation is a posttranslational modification which is mediated in cells by ADP-ribosyltransferases (ARTs) of the ARTD family, which add ADP-ribose (ADPr) to their targets while releasing nicotinamide from co-factor NAD^+^ (1, 2). Best studied is the modification with chains of ADPr, termed poly(ADP-ribosyl)ation (PARylation), which amongst other processes has a demonstrated role in the DNA damage response, regulation of protein stability and Wnt signalling. Inhibitors of the enzymes capable of generating PAR-chains, the poly-ARTs, have found clinical applications in specific breast and ovarian cancers, with more clinical trials underway (3).

Much more enigmatic is its little sibling, the modification of proteins with just one ADP-ribose moiety (MARylation). The ARTD family contains 17 members (4, 5), of which only four are capable of generating chains of ADP-ribose, namely PARP1, PARP2, TNKS1 and TNKS2. The other members of this protein family have mono(ADP-ribosyl)transferase activity, or no proven activity yet in the case of PARP13 (6, 7). Despite the potentially misleading name PARP for MARylating enzymes, this term was kept for historical reasons (8). MARylation may play a role in immunity and transcription amongst other functions (2, 9-12), which are very varied: PARP10 may regulate kinase activity (13-15) and replication (16), PARP12 localises to the Golgi and stress granules and may function there (17, 18), PARP14 was reported to act as transcriptional co-activator (19), but also to be present at focal adhesions (20) and involved in DNA replication and repair (21, 22). Recent work on PARP7 points to a role in both immunity, regulation of the cytoskeleton and suggests that it may serve as anti-cancer drug target (23-25). These highly diverse functions of MARylation and associated transferases have been reviewed in detail (2, 10-12).

Different amino acids were identified as ADPr acceptors in recent years using different approaches including mass spectrometry: glutamates, serines, tyrosines, histidines and most recently cysteines (24, 26-29). In addition to the modification of proteins with ADPr, recent *in vitro* data indicate that some of the mammalian PARPs can also modify nucleic acids as summarised elsewhere (30, 31), although it is not clear yet what role RNA and DNA MARylation has in cells.

MARylation of both proteins and nucleic acids is a reversible modification, with different hydrolases removing the modification from different acceptor sites. ARH1 removes ADPr from arginine (32), ARH3 from serine (33) and MACROD1/MACROD2/TARG1 reverse the modification of acidic residues (13, 34-36). PARG and ARH3 both cleave the *O*-glycosidic bond between ribose moieties in PAR and are thus responsible for reversing PARylation (36). PARG leaves the terminal ADP-ribose on the substrates, whereas ARH3 can remove this if linked to serine. No enzyme has been identified yet that reverses the ADP-ribosylation of cysteine residues. Not only mammalian genomes encode for macrodomain-containing proteins, they are also present in other organisms (36). Triggered by the COVID19 pandemic, the spotlight has recently been on the SARS-CoV-2 macrodomain protein Mac1, which was shown to be an ADP-ribosylhydrolase with thus far unknown substrates (9, 37, 38). Not only certain transferases, but also a number of the mammalian hydrolases has been suggested to drive certain aspects of tumorigenesis such as transformation, growth and invasiveness (39, 40). This has been difficult to verify, as antibodies for the hydrolases were poorly characterised and antibodies specific for MARylation were not available at all.

Research in the area of MARylation has been held back by lack of tools for the analysis and detection of modified substrates. In recent years two major breakthroughs have occurred: optimised mass spectrometry methods, which are now reliably able to detect the modified proteins present in cells (29, 41, 42), and the development of multiple antibodies and other reagents which detect this modification. This provides a great opportunity which has to be enjoyed with caution: most of the reported detection tools were tested on specific substrates, but their actual epitopes have been poorly mapped. It is thus not clear whether it is possible to directly compare one study performed with one antibody to another study using another detection tool. The first specialised readers of MARylation are the macrodomains of PARP14, which have been used to detect intracellular MARylation using live-cell imaging (43, 44). This was turned into a commercially available detection reagent by fusion of specific macrodomains to the Fc region of rabbit immunoglobulin (45). Next an antibody was generated against a peptide with a MARylated lysine which appears to detect MARylation in cells efficiently as well as PARylation (46). This was followed by the creation of a modified version of the MAR-hydrolase Af1521, which has increased binding affinity compared to the wildtype protein but is still catalytically active (29). Next, antibodies were generated against MARylated serines as well as a general ADP-ribose antibody (47). Most recently, a polyclonal antibody raised against MARylated peptides was raised in rabbits for detection of MARylation (48). Several reagents are thus available that can be utilised to study the *in vivo* function of MARylation, however, none of these reagents have been compared with each other. It is not clear whether there are differences in substrate recognition, whether some may preferentially bind to specific MARylated amino acids or even recognise part of the protein backbone. It is highly probable that the MAR-specific reagents do recognise either the specific ADPr-protein bond or part of the protein surrounding, as otherwise they would be expected to be efficient tools to detect PARylation as well.

We have compared the above-mentioned different reagents to map their respective specificities. For this purpose, we enzymatically generated protein substrates MARylated on specific amino acids, such as serine, arginine and glutamate, or peptides chemically modified on serine, cysteine and threonine, as well as MARylated nucleic acid substrates, and tested all the detection reagents for ADP-ribosylation that were available to us. We verified the specific modification of our substrates by using a panel of hydrolases, which reverse the modification only from specific targets as expected. We have included a detection reagent based on murine PARP14 macrodomains, which was previously described to have higher affinity for ADPr than the human macrodomains (43) and appears to be an efficient tool for immunoprecipitation of ADP-ribosylated nucleic acids. After determining the specificities of the antibodies on *in vitro* generated substrates, we next asked whether differences exist in their recognition of the modification introduced by different PARPs overexpressed in cells and tested their suitability for immunofluorescence using different fixation methods. To be able to do this, we optimised the preparation of cell lysates to ensure maximum retention of the ADPr signal. Collectively, our work deciphers which reagents are suitable for which purpose and has optimised sample preparation procedures which allow detection of low ADP-ribosylation levels.

## RESULTS AND DISCUSSION

### ADPr-detection reagents have different specificities *in vitro*

To be able to compare the currently available ADP-ribosylation detection reagents, we required suitable, well-defined substrates. Therefore, we assembled a collection of ADP-ribosyltransferases with known substrate specificity. Using these enzymes, we can generate proteins specifically modified on cysteine by pertussis toxin (PT), acidic amino acids by PARP10, arginine by mART2.2 and SpvB, serine by PARP1/HPF1 or PAR by PARP1. Recent studies reported modification of other amino acids, such as histidine or tyrosine, however, the respective enzymes are not yet known and could therefore not be included. The majority of proteins employed here were purified from bacterial expression systems, with the exception of PARP1, which was immunoprecipitated using a GFP-tag from HEK293T cells (**Figure 1a**). All purified proteins are active and either automodify or ADP-ribosylate a specific target (**Supplementary Figure 1**), with the exception of the arginine-ART mART2.2, which modifies many proteins present in a cytosolic extract. Nuclei and mitochondria were removed as they contain largest amounts of PARP1, TARG1 and MACROD1 and could confound the ADP-ribosylation assay. We noticed automodification also for pertussis toxin, which is in line with previous observations using this truncated version of the protein (49). To confirm the specificity of the ADP-ribosylation reactions, we incubated these substrates with a panel of hydrolases. For PARP10 and PARP1/HPF1, we made use of specific inhibitors to stop the reactions before adding hydrolases (50). ARH1 is only capable of reversing arginine modification, whereas ARH3 hydrolyses both PAR as well as removes ADP-ribose linked to serine. PARG is only capable of hydrolysing the glycosidic bond between ADP-ribose moieties and should not be able to remove the ADP-ribose linked to the proteins. MACROD1, MACROD2 and TARG1 have been described to reverse the modification of acidic residues (13, 34). For the cysteine-linked ADP-ribosylation introduced by PT no erasers are known to date. We purified the known erasers of ADP-ribosylation from bacteria: MACROD1, MACROD2, TARG1, PARG, ARH1 and ARH3 (**Figure 1a**). The majority of described activities we could confirm, for example MACROD1, MACROD2 and TARG1 reverse the modification introduced by PARP10 although not fully, as has been seen before (**Figure 1b and Supplementary Figure 2**) (6, 51). We could furthermore confirm that ARH3 reverses both PAR and serine-linked MARylation (**Figure 1c**), but also appears to have some activity towards PARP10 (**Figure 1b**),. This was hinted at in earlier work studying reversal of ADP-ribosylation by PARP10 (6) and would be expected if PARP10 also automodifies on serine as reported (51). ARH1 reverses the arginine modification introduced by mART2.2 very efficiently, but has no activity towards other modified amino acids (**Figure 1d**). As the hydrolases reverse the diverse modifications as expected, we concluded that our *in vitro* substrates are modified on the expected amino acids. None of the hydrolases were able to reverse the modification from cysteine as generated by pertussis toxin (**Figure 1e**). This raises the possibility that additional mammalian intracellular hydrolases remain to be discovered. Alternatively, the automodification of the truncated pertussis toxin we used may not represent an accessible substrate for the hydrolases tested and we can thus not exclude that they are capable of reversing cysteine MARylation on different substrates.

**Figure 1.**
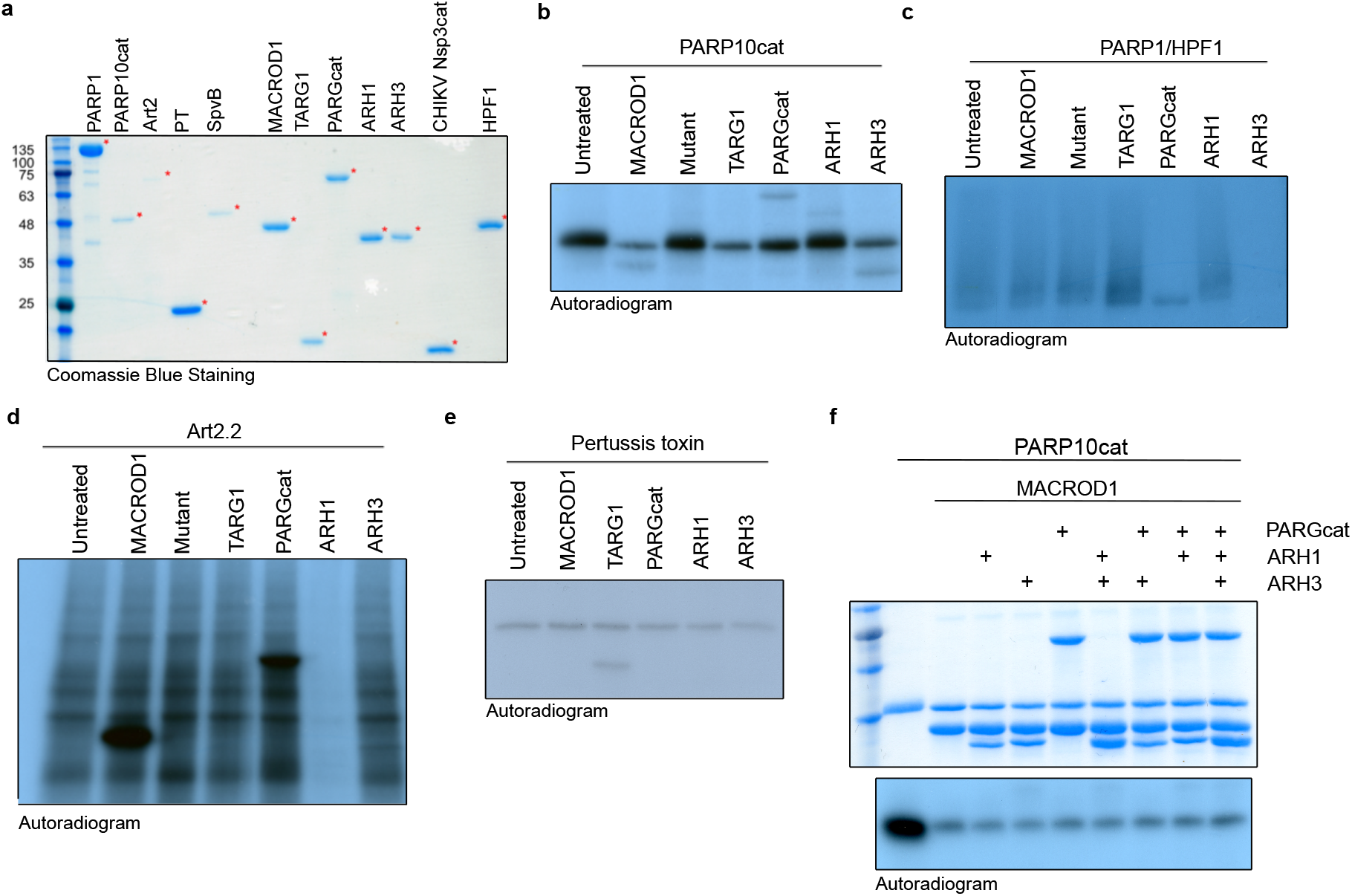
Enzymatic generation of specific ADP-ribosylated substrates. **(a)** Coomassie blue staining of the purified transferases, hydrolases and co-factors used in this study. (**b-e**) Indicated transferases were incubated with ^32^P-NAD^+^ and incubated for 30 minutes at 37°C. A cytoplasmic extract was provided to supply the Art2 and pertussis toxin substrates. After the transferase reaction, OLU35 was added to inhibit PARP10 or olaparib to PARP1/HPF1, followed by a 30 minute hydrolase reaction. Samples were run on SDS-PAGE and incorporated radioactivity was visualized using film. (**f**) PARP10 was automodified using ^32^P-NAD^+^, the reaction was stopped with OLU35 and indicated hydrolases were added. The Coomassie staining is displayed to show the relative amounts of hydrolases added. The samples were analysed as in (b).

In addition, we also noted that none of the hydrolases is able to completely reverse PARP10 modification and speculated that PARP10 may be promiscuous and able to modify more than one type of amino acid. Partial reversal of PARP10 automodification by PARG and ARH3 was observed before, which may be reversal of oligomers that PARP10 was reported to generate (6). A more recent study using recombinant proteins also hinted at PARP10 promiscuity, as the modification of serine, arginine and glutamate by PARP10 was detected using mass spectrometry (51). We incubated the protein with different hydrolases and combinations thereof. We were not able to further decrease the modification after removal of the majority of the signal by MACROD1, as no other hydrolase leads to further decrease of the remaining signal (**Figure 1f**). Three potential explanations come to mind: first, it in addition modifies an amino acid for which we have not identified a hydrolase yet, such as cysteine, second, it is an artefact of the *in vitro* reaction and for example on the terminal amine, or third, the bond to the side chain of an acidic amino acid migrates from the C1’ to the C2’ or C3’ position and thus is no longer available to known hydrolases.

Having thus confirmed the specific modification of our enzymatically generated substrates, we next performed western blots to test the specificities of the antibodies and detection reagents available to us. We generated large amount of substrate (**Supplementary figure 1**), stored this at -20°C and proceeded with western blots once radioactivity decayed with the detection reagents described in **Table 4**. We first used the macrodomain-based detection reagents on these substrates, Reagents I-III, which are based on either human or murine PARP14 macrodomains or contain the aforementioned Af1421 fused to an Fc (**Figure 2a-d**). We could confirm their reported specificity for MARylation over PARylation, as also for example the PARP1-HPF1 sample resulted in a more specific band instead of a smear. We developed the murine PARP14 macrodomain-based detection reagent, Reagent III or mPARP14-m2m3-Fc, as affinity for ADPr was reported to be higher for the mouse than human macrodomain proteins (43, 44). We also generated a control, Reagent IIIm or mPARP14-m2m3-GE-Fc, which has impaired ADPr binding. This mutant does not show these specific signals. The polyclonal antibodies Reagent IV (**Figure 2e**) and V (**Figure 2f**) have very similar properties on these *in vitro* substrates. The pan-ADPr antibody Reagent VI shows highest affinity for Arg-ADPr and Ser-ADPr (**Figure 2g**). We also tested an antibody which was described to detect PARylation, Reagent VII. It recognises PARylation as stated, but also for MARylation bands are detectable (**Figure 2h**), which raises the question whether a fraction of the modification generated by these enzymes are oligomers, or whether the antibody can detect single ADPr moieties. It for example detects the modification introduced by PARP10, which could reflect a minor oligo-ADPr-transferase activity which PARP10 was reported to possess (6). A significant background is present, despite identical processing of the blots, making this antibody potentially less suitable. Interestingly, the polymers are hardly detected by any of the antibodies in this experiment, although some in theory should. It is possible that the long storage in the freezer led to degradation of these samples. We therefore performed new analyses of GFP-PARP1 immunoprecipitated from HEK293T cells. In parallel we performed an experiment using radioactively labelled PAR chains, demonstrating effective reversal of the oligomers by PARG with exception of the last ADPr, whose resistance to reversal by MACROD1 confirms the specificity of this signal: oligo-ADPr attached to the substrate via serine (**Supplementary Figure 3**). Reagents I and VI show weakest staining overall, the other tested reagents all detect PARylation as well as Ser-ADPr efficiently with the exception of Reagent III, which does not detect PARylation at all and Ser-ADPr relatively weakly (**Figure 2h**). To be able to test some substrates we cannot generate enzymatically, we next slot-blotted chemically synthesised peptides modified with ADPr on serine, cysteine and threonine (52), including a peptide modified with phospho-ribose as negative control, and tested the affinity of the different reagents towards these substrates (**Figure 2i**). The results are consistent with previous observations: the Reagent III and VII lead to high background staining (**Supplementary Figure 4)**; the anti-ADPr antibodies Reagent IV and V detect cysteine, threonine and serine ADP-ribosylation, as does the reagent based on Af1521, Reagent II. The fact that both Reagent I and Reagent III, which are based on human and mouse PARP14 macrodomains respectively, do not recognise any of these peptides implies that they do recognise part of the protein surrounding the modification, which may not be present in the tested peptides. This offers a tentative explanation why these modules are specific for MARylated proteins: if they would detect ADPr or a part thereof, they should at least detect the terminal ADPr of PAR chains as well. Binding of the module to the surrounding backbone would explain their exclusive detection of MAR-but not PARylated substrates.

**Figure 2.**
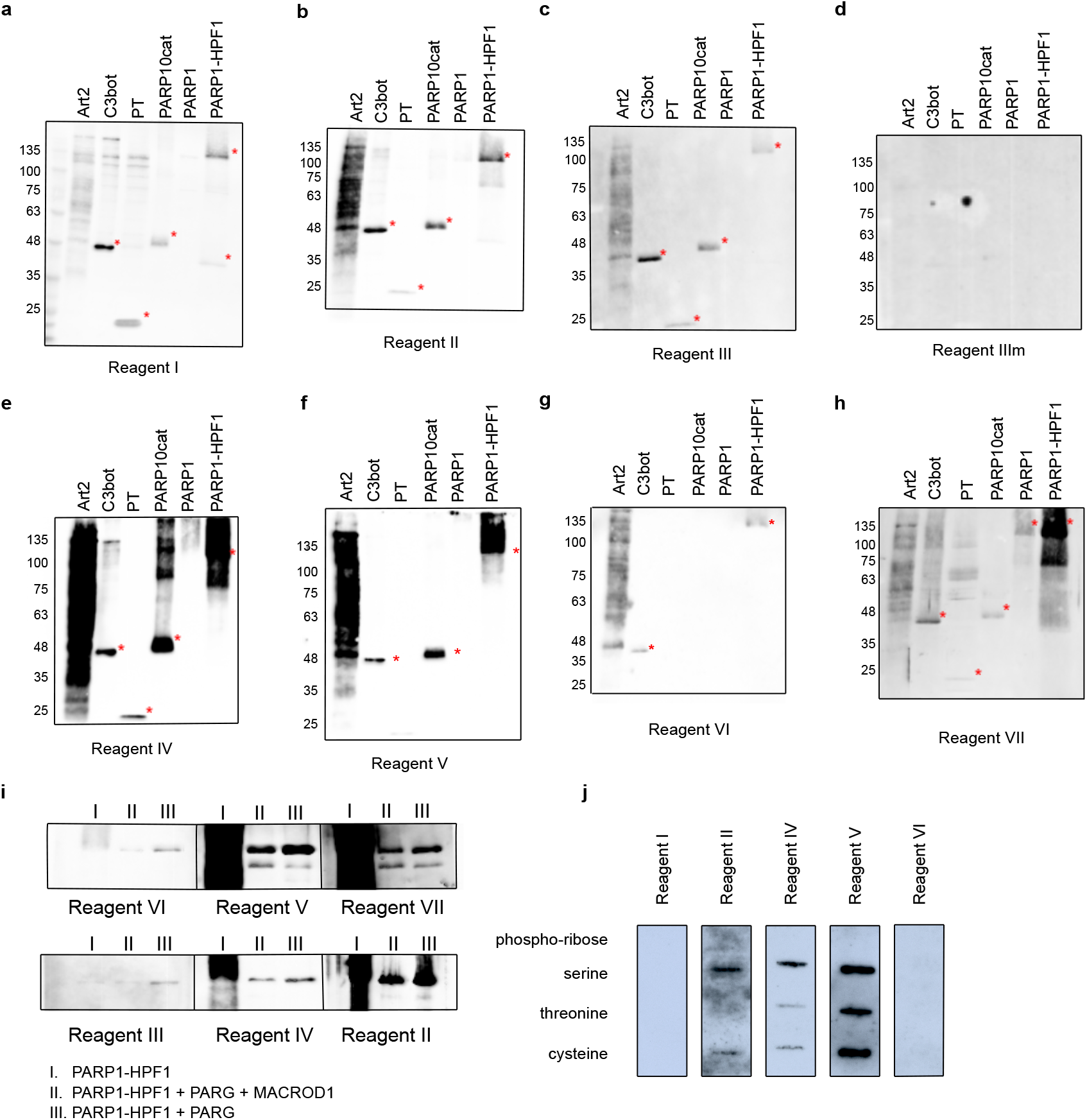
Specificity of the reagents towards *in vitro* modified substrates. (**a-h**) ADP-ribosylation reactions from 1b were loaded on multiple gels and blotted. Membranes were blocked in 5%-milk in TBST and incubated with primary antibodies overnight. (**i**) GFP-PARP1/His-HPF1 was incubated with NAD^+^ (I), followed by PARG and MACROD1 (II) or PARG (III) treatment. After blotting, modification was detected the same reagents (**j**) 2µM chemically synthesised peptide modified with either phospho-ribose or ADP-ribose on serine, threonine or cysteine were slot blotted and analysed using the indicated reagents. Asterisks indicate the transferases.

### Addition of olaparib during lysis and avoiding boiling of samples are essential for detection of endogenous ADP-ribosylation states

After having determined their specificity on *in vitro* substrates, we next determined which signals the antibodies detect in HEK293T lysates. We overexpressed almost all full-length GFP-tagged PARPs, as well as GFP alone, and first confirmed expression of these proteins (**Figure 3a**). PARP2 was not included due to its expected redundancy with PARP1 and for PARP14 we were not able to generate a full-length construct containing an intact N-terminus. Overexpression of these enzymes led to degradation of some enzymes, apparent from the smaller products visible in the anti-GFP blot. We nevertheless analysed the MARylation signal using Reagent V. We used this reagent to optimise experimental conditions, as it gives robust signals on the *in vitro* substrates and is commercially available in large amounts. We observed a relatively equal signal in most lanes, including the GFP-transfected control lane (**Figure 3b**). PARP1 is highly abundant and activated by damaged DNA, which is available in high abundance during our RIPA-based lysis. To avoid false PAR/MAR signals from PARylation introduced by PARP1 during cell lysis, we performed all subsequent experiments with addition of the PARP inhibitor olaparib to the lysis buffer. When preventing PARylation during lysis, the pattern detected by the antibody changes dramatically (**Figure 3c**): no activity can be seen for most of the poly-ARTs, with the exception of TNKS2, which could be partially caused by omitting a PARG inhibitor during lysis. A signal is present for the majority of mono-ARTs. PARP6, -7, -8, -10 and -15 overexpression leads to modification of diverse proteins, whereas for PARP11 and PARP12 only few substrates were detected. As it was shown that inhibition of the proteasomal degradation system may lead to an increase of overexpressed PARP7 MARylation (46), we tested whether inhibition of the proteasome leads to an increase of the signal in our experiments, to confirm that the observed signal is ADP-ribosylation. For most enzymes we observe only slight differences in the MARylation signal after proteasome inhibition, however, for PARP6, PARP7, PARP12 and PARP15, the signal appeared stronger upon proteasomal inhibition (**Figure 3c**). This may imply that MARylation is a mark that can, directly or indirectly, lead to the target proteins’ degradation. For others proteins, like PARP12, only distinct bands, which may correspond to the overexpressed proteins themselves, are increased upon MG132 treatment. It is possible that the MARylation signal present upon overexpression of individual PARPs is not derived from modification introduced by this PARP, but from another PARP which became activated upon introduction of the exogenous enzyme. There are examples of PARPs modifying one another, as PARP7 for example is capable of modifying PARP13 (24). When lysing cells under basal conditions, only a low signal for endogenous MARylation is present, and this is not increased upon transfection of for example a GFP-encoding plasmid, indicating that it is not the transfection stress which activates the endogenous ARTs. We also noted that western blots were on occasion not reproducible when performed only a few days apart using the same lysates, which prompted us to test the effect of different sample preparation conditions on western blot outcome. For these analyses, we re-used the samples from figure 3c and analysed the enzymes which rendered the highest MARylation signal. When samples were frozen, boiled for 5 minutes in SDS-sample buffer and analysed on gel again, the signal-to-noise ratio was worse, implying an increase of unspecific modification or a loss of specific modification upon freezing and boiling of the samples (**Figure 3d-e**). When loading the same frozen samples without boiling, a few specific bands seem to be preserved which were lost upon boiling, for example in the PARP11 and PARP12 samples (**Figure 3d**), however also here the signal-to-noise ratio is worse than it was in the initial analysis (**Figure 3c**). Previous work has also demonstrated thermolability of the cysteine ADP-ribosylation of GAP-43, as approximately 50% of the modification is lost upon incubation for 10 minutes at 95°C (53). We utilised the same peptides as well as the automodified PARP10 catalytic domain, and slot-blotted them untreated, or heated at different temperatures to test whether the modification is thermolabile (**Figure 3f and Supplementary Figure 5**). The modification of the peptides and PARP10 decreases with incubation time and temperature. At 95°C the modification is lost rapidly from all peptides as well as from PARP10 itself, whereas at 60°C the modification appears more stable although also diminishes with longer incubation times. It is possible that also on other amino acids the modification is thermolabile. In subsequent experiments we always analysed fresh lysates, prepared with olaparib and only briefly heated at 60°C.

**Figure 3.**
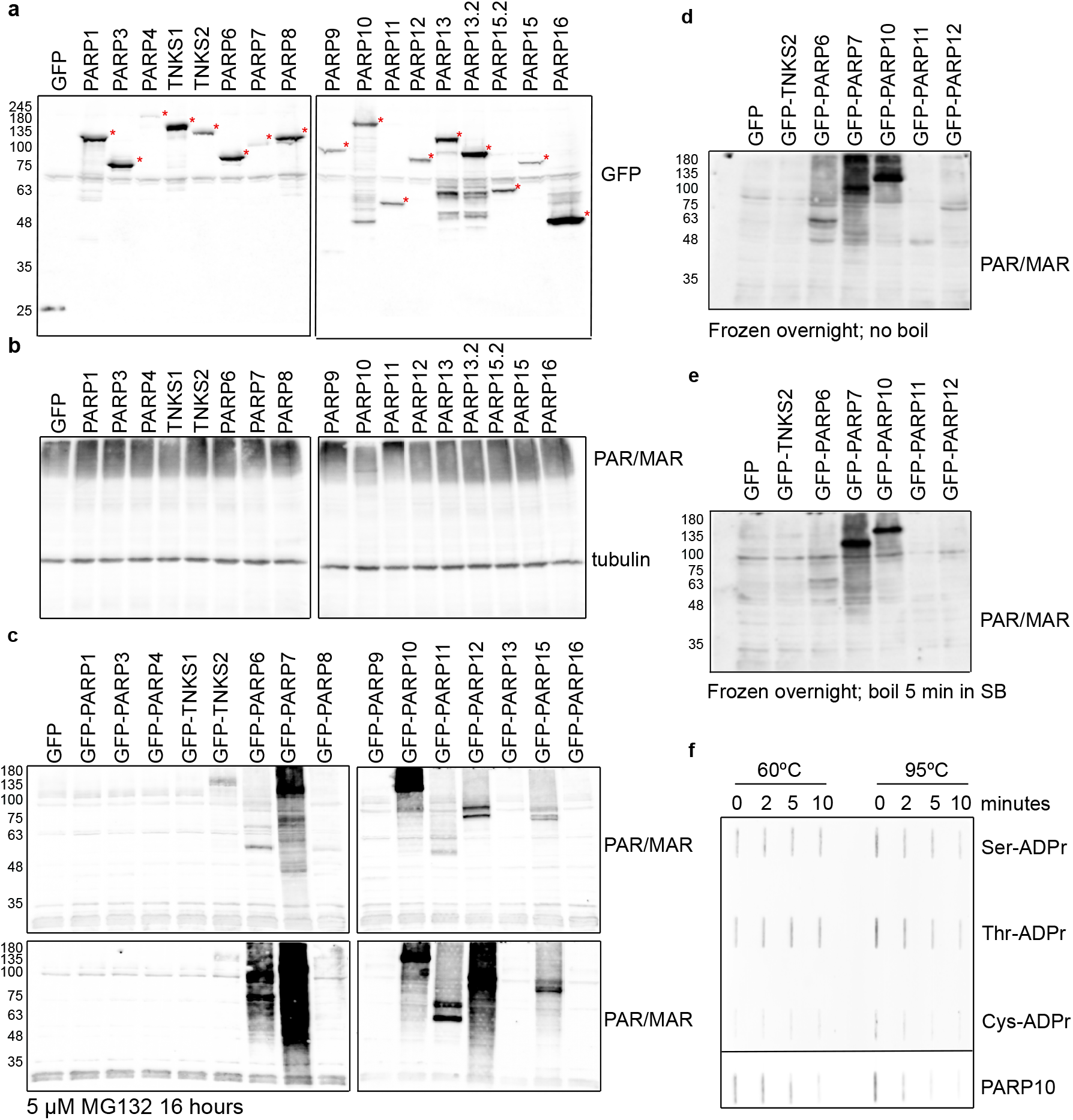
Optimised lysis conditions are required to prevent ADP-ribosylation occurring during lysis or degradation during sample preparation. (**a**) HEK293T cells were transfected with indicated GFP-PARPs, lysed in RIPA buffer and analysed using western blotting with a GFP antibody (**b**) Same lysates as in (a), but analysed with an ADPr antibody Reagent V. (**c**) HEK293T cells were transfected with indicated GFP-PARPs and lysed in RIPA supplemented with olaparib or in addition treated with proteasome inhibitor MG132 before lysis. Western blots were analysed with an ADPr antibody Reagent V. (**d-e**) Untreated lysates from (c) were frozen at -20°C and heated at (d) 60°C or (e) 95°C before loading the SDS-PAGE. Resulting western blots were analysed with an ADPr antibody Reagent V. (**f**) ADP-ribosylated peptides and automodified PARP10 catalytic domain were slot-blotted untreated, or heated at 60°C and 95°C for 2, 5 or 10 minutes. The blot was analysed using an ADPr antibody Reagent V. The PARP10 blot was exposed shorter due to the stronger signal, different exposures are provided in the supplements.

### Not all reagents perform equally well on endogenous ADP-ribosylation

Having established the conditions best suited for lysis, we next analysed ADP-ribosyltransferases which modify different targets: TNKS2 was included as poly-ART, PARP7 as cysteine-modifying enzyme, murine ART2.2 for arginine, PARP10 for its postulated major modification of glutamate and PARP6 as example of ART for which specificity has not been determined yet. The antibodies Reagent IV and V show a robust modification signals for most enzymes (**Figure 4a-b**). In general, the antibody Reagents IV-VI detect the arginine modification introduced by mART2.2 well (**Figure 4a-c**). In contrast to these, a TNKS2 signal is present when using anti-PAR Reagent VII as expected (**Figure 4d**). Reagent II detects most substrates, both polymers from TNKS2 as well as MARylation from the different enzymes (**Figure 4e**). Reagent I detects MARylation introduced by PARP6, PARP7 and PARP10 (**Figure 4f**). The weakest but perhaps most specific signal is visible when the blots are probed with Reagent III (**Figure 4g**), as here only PARP7 and PARP10 lead to clear signals. These signals are absent with the binding-deficient mutant Reagent IIIm (**Figure 4h**). Expression of the GFP-constructs was determined using an anti-GFP western blot (**Figure 4i**). Reagent I, which is based on the human macrodomains from PARP14, in addition detects arginine. The greatest variability is visible for PARP6: some reagents detect no signal at all, such as reagent I and VI, where others detect robust MARylation. This highlights the necessity of ongoing reagent evaluation, as it appears that the suitability of a reagent is dependent on the enzyme of interest.

**Figure 4.**
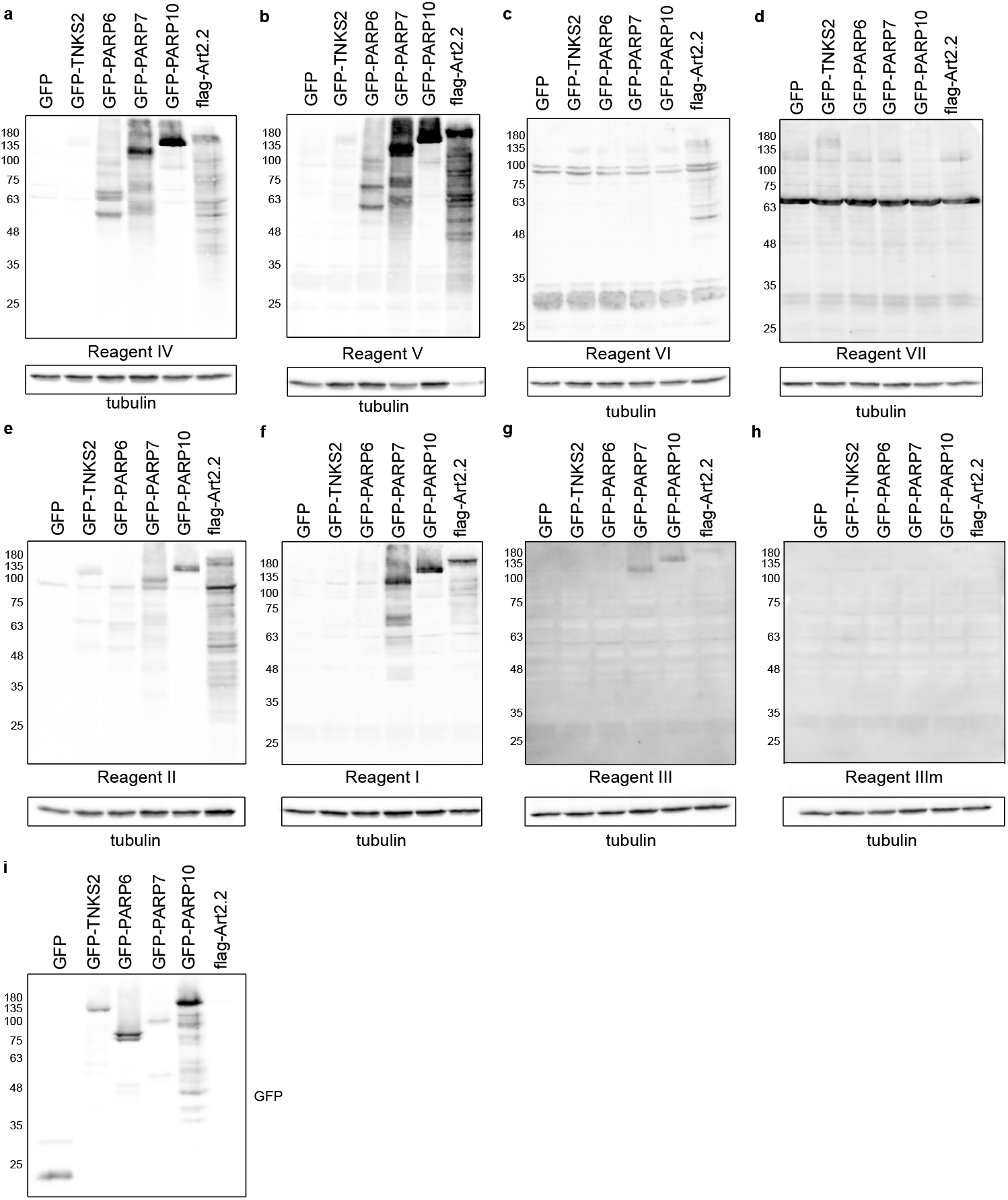
The anti-ADP-ribose detection reagents have diffferent specifities for substrates modified in cells. (**a-h**) HEK293T cells were transfected with the indicated GFP-tagged PARP constructs, GFP as control or flag-tagged murine ART2.2. 24 hours after transfection, cells were lysed in RIPA buffer supplemented with olaparib and analysed using SDS PAGE. The western blot was detected with the indicated reagents. (**i**) Western blot showing the transfection levels of the GFP-transferases.

### Only some reagents are suitable to detect RNA ADP-ribosylation

As recent publications have shown that also nucleic acids can be MARylated at least *in vitro*, we next determined whether these different reagents can be used to detect the modified nucleic acids. We generated MARylated RNA oligonucleotides and confirmed their modification using denaturing urea-PAGE and SYBR gold staining to visualise the nucleic acids in gel (**Figure 5a**). We purified the different RNA species and slot-blotted them alongside automodified PARP10 as control (**Figure 5b**). Several reagents can be used to detected MARylated RNA: both the antibodies Reagent IV and V, as well as the macrodomain-based Reagent II and III detects the modification. We performed a similar experiment with NAD^+^-capped RNA, which we produced as described before (54). To test the procedure, we generated RNA with a radiolabelled NAD^+^-cap and subjected this to DXO treatment, which degrades specifically NAD^+^-capped RNAs (54) (**Figure 5c**). None of the antibodies tested can be used to detect the NAD^+^-cap (**Figure 5d**), which agrees with dot-blots where NAD^+^ could not be detected by the anti-ADPr antibody (48). Adenylylation is another modification which closely resembles ADP-ribosylation and may be recognised by these reagents. We generated 5’-adenylated ssRNA and slot-blotted these alongside ADP-ribosylated RNA and noted that Reagents I, V and IV can detect the modification, whereas the macrodomain-based reagents II and III are not able to bind this. (**Supplementary Figure 6**). This further confirms that the reagents recognise significantly different epitopes. Lastly, we enriched *in vitro* ADP-ribosylated ssRNA using GFP-mPARP14-m2m3-Fc wildtype or GE mutant. mPARP14-m2m3 efficiently binds the modified ssRNA, but not the unmodified RNA (**Figure 5f**). The binding-mutant does not interact with either modified or non-modified RNA. In this pull-down, an additional band is enriched with m2m3 wildtype, which could imply that PARP10cat can generate an oligomer, which the module preferentially binds, or that it modifies multiple sites on each RNA, also leading to enhanced precipitation. In addition to their usage to study MARylated proteins, a subset of the available antibodies and detection reagents can thus be used to start studying the presence and function of nucleic acid ADP-ribosylation in cells. The macrodomain-based reagents are ideal to identify the modified RNAs in cells, as the binding mutants can be used to monitor non-specific binding to both module and column. These data also highlight that their epitopes are diverse, as not all function equally well on nucleic acid substrates.

**Figure 5.**
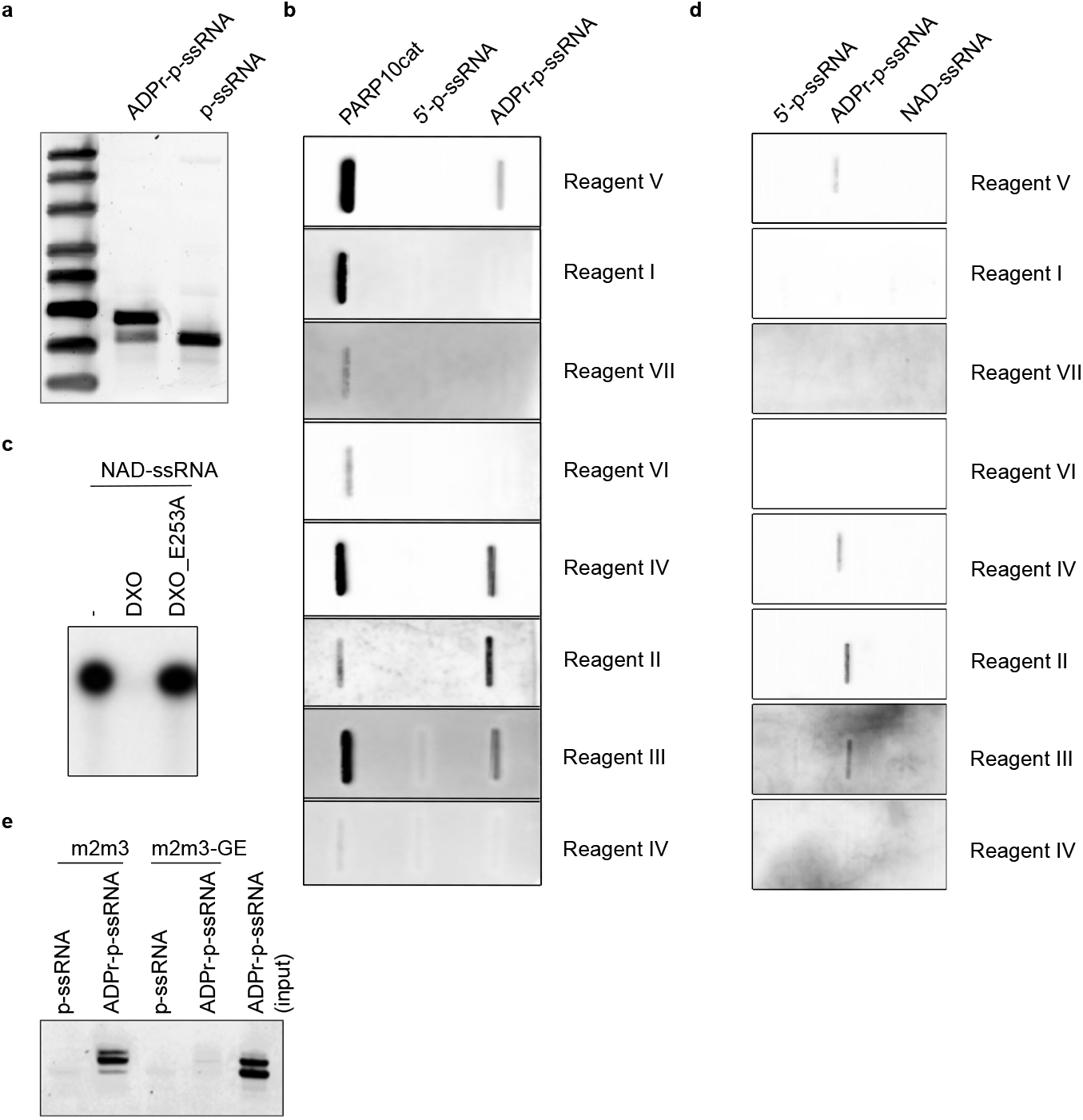
A subset of the ADP-ribosylation reagents can be used to detect and enrich MARylated ssRNA but not NAD^+^-capped RNA. (**a**) A phosphorylated, single stranded RNA (p-ssRNA) was ADP-ribosylated with the catalytic domain of PARP10 ranging from amino acids N818-T1025 (PARP10cat), resolved on urea-PAGE and analysed using SYBR gold staining. (**b**) A slot blot was performed with automodified PARP10cat, unmodified ssRNA or ADP-ribosylated ssRNA. The MARylated ssRNA was treated with proteinase K and purified before slot-blotting. The blot was analysed using the indicated detection reagents. (**c**) The product of an *in vitro* transcription reaction with ^32^P-labelled NAD^+^ was analysed using urea-PAGE and autoradiography. DXO specifically removes an NAD^+^-cap and degrades the RNA oligo, whereas an inactive mutant is unable to remove the NAD^+^-cap. (**d**) As in b, but now with NAD^+^-capped RNAs. NAD^+^-capped RNAs were generated during an *in vitro* transcription reaction, treated with proteinase K and purified before analysis by slot-blotting. (**e**) ssRNA or ADPr-ssRNA generated as in (a) were incubated with magnetic agarose beads coated with the indicated mPARP14-m2m3 constructs. After extensive washing, samples were eluted from the beads and analysed on urea-PAGE

### Immunofluorescence staining of ADP-ribosylation is highly dependent on fixation method

Lastly, we tested the reagents’ suitability for immunofluorescence staining and confocal microscopy. We fixed cells using cross-linking agents paraformaldehyde (PFA) and glyoxal, as well as the dehydrating agent methanol. The CST antibody, Reagent V, gives the strongest signal: if identical laser strengths are used for the other reagents, the signals are very low (**Figure 6a** and **Supplementary Figure 7**). The relatively weak staining obtained with Reagent II, based on Af1521, could be caused by its catalytic activity during our procedures at room temperature, which should be restricted during our incubations for western blot at 4°C. Using PFA as fixative, we can confirm the cytoplasmic staining reported before with Reagent V (**Figure. 6a**) (48, 55). Glyoxal fixation in general resembles the methanol fixed samples and leads to low staining intensity with most reagents, but gives rise to nucleolar staining with the CST antibody (**Figure 6a**). Upon H2O2 treatment, the ADP-ribosylation signal becomes strongest in the nucleus (**Figure 6b**). The discrepancy between cytoplasmic stainings in untreated cells fixed with PFA can have several possible explanations: either the respective epitopes are not exposed after fixing with methanol, or something is stained which is washed out by methanol but crosslinked by PFA, such as small metabolites. We expected similar staining patterns between PFA and glyoxal, as both are cross-linking agents. One key difference, however, is that glyoxal does not crosslink RNA as well, in contrast to PFA. In theory, it is possible that the strong cytoplasmic signal observed with a number of the reagents is mitochondrial RNA. Both antibodies which efficiently recognise slot-blotted MARylated ssRNA, also stain cytoplasmic structures after PFA fixation but not after glyoxal. We incubated the slides with either RNAse or DNAse to remove signals derived from potentially MARylated RNA or DNA, neutral hydroxylamine to reverse modification of acidic residues or mercury chloride to reverse modification of cysteines. All treatments lead to some reduction of the signal, however, the strongest change is visible in the hydroxylamine treated samples (**Figure 6c**). This implies that the majority of detected modification in mitochondria is a protein ADP-ribosylated on acidic residues, which leaves the question open why PFA and glyoxal stainings are different. Alternatively, the hydroxylamine treatment may interfere with the antibody epitope and therefore lead to loss of staining. It can thus not be concluded with certainty whether DNA, RNA or proteins are the main source of ADP-ribosylation staining in mitochondria.

**Figure 6.**
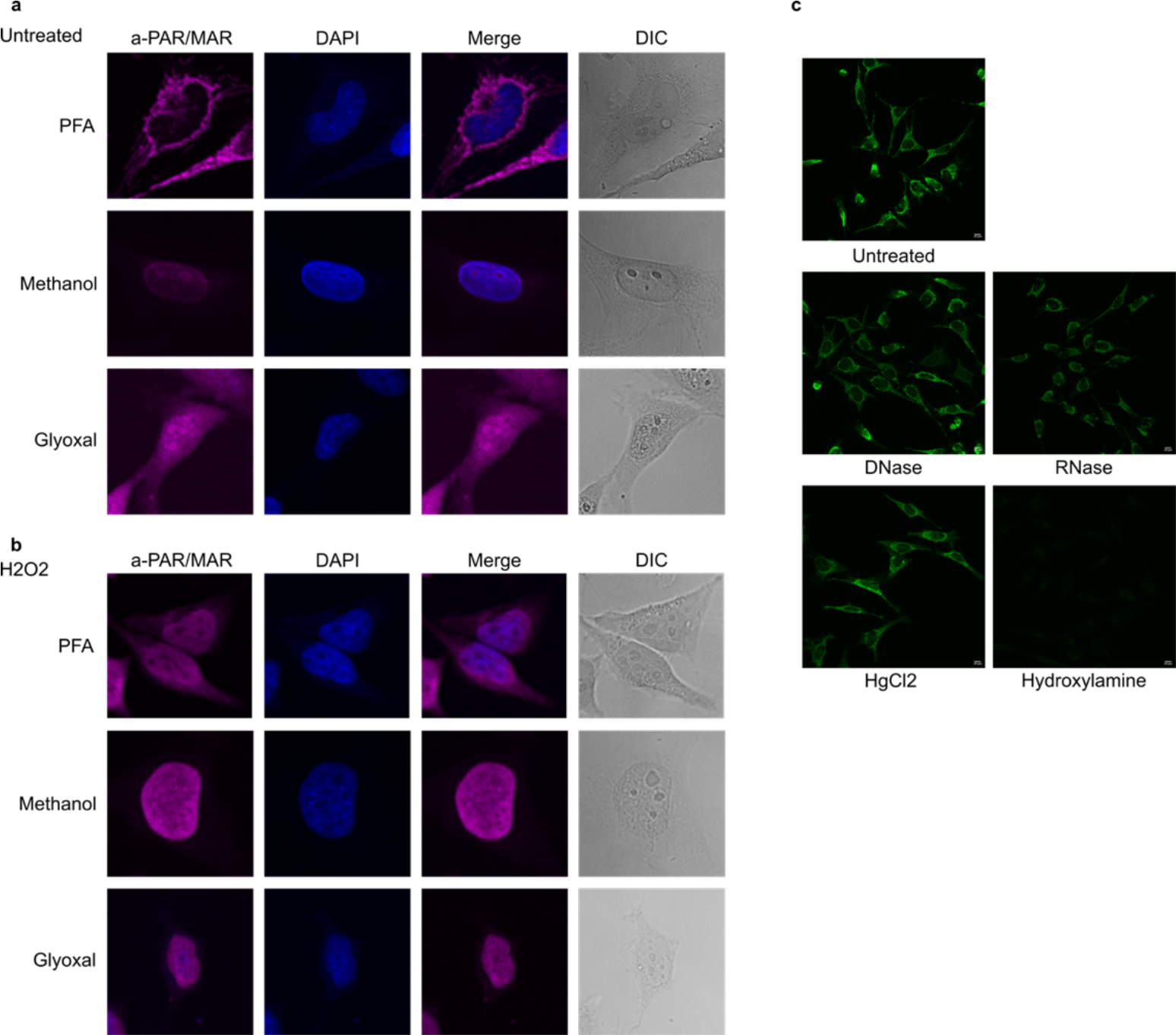
ADP-ribosylation detection reagents stain different structures depending on fixation method used. (**a**) HeLa cells were seeded on glass coverslips and fixed using paraformaldehyde (PFA), ice-cold methanol or glyoxal. Reagent V was used to stain the cells and DAPI was applied to stain the nuclei. (**b**) As in a, but before fixation cells were treated with 0.5 mM H2O2 for 10 minutes. All images were taken using a confocal microscope with identical laser intensity and settings across all samples. (**c)** HeLa cells were seeded as in (a) and fixed using PFA. Fixed samples were treated with either DNase or RNase, HgCl2 or hydroxylamine. ADP-ribosylation was visualised using the same PAR/MAR antibody as in (a) and images taken using confocal microscopy at the same settings for each sample.

## CONCLUSIONS

In this work, we set out to compare the affinities of the different reagents available to detect MARylation. They have different strengths and weaknesses: many also recognise PAR, making them less suitable to study exclusively MARylation. One other aspect which may interfere with the blots performed with these reagents, is the existence of AMPylation as protein modification (56): those reagents which are independent of amino acid linkage and surrounding amino acids, may also detect proteins modified with AMP, as was demonstrated for a pan-ADPr reagent (57). The macrodomain-based reagents show higher specificity towards MARylated substrates, but in general bind weaker to the modified proteins. A major advantage of the macrodomain-based reagents, is that these can be generated any time, thus avoiding the risk that reagents will become unavailable as was the case with the anti-PAR antibody from Trevigen used here, which is not available anymore. A further advantage is the possibility to create ADP-ribose binding mutants, which can be used as negative control to eliminate the proteins which bind unspecifically. Despite weaker signals in western blots, these macrodomains and their mutants may prove very useful to immunoprecipitate MARylated proteins and especially RNAs with high specificity.

As the concentration of the reagents provided by other researchers was partially unknown, we cannot determine their absolute affinities, but merely compare which reagent is best suited for the respective substrates. It should also be kept in mind that production methods are different, ranging from purification from sera for polyclonal antibodies and hybridoma culture for monoclonal antibodies, to the supernatant of cultured cells for eAf1521 to *E*.*coli* for mPARP14-m2m3-Fc. These differences could potentially influence the outcomes as well and may require further optimisation, especially for mPARP14-m2m3-Fc, where it is feasible to reduce the background noise by further optimisations of purification and storage conditions. Not only is the use of the correct detection reagent for specific substrates important, we also illustrated how sample preparation can influence the apparent ADP-ribosylation signal. Routine boiling of western blot samples may have detrimental effects and lead to false-negative results. Inhibition of PARP1 is necessary to prevent induction of poly(ADP-ribosyl)ation during lysis, which may also be true for other ARTD family members as well as the reverse hydrolase reaction. Further ART and hydrolase inhibitors should be added during lysis or even to the cell cultures, if available. Similarly, the fixative chosen for immunofluorescence approaches influences the structures stained by the different reagents. Sample processing can thus hugely influence experimental outcome and ought to be both considered and documented carefully.

An interesting question arising from the immunofluorescence images is why the mitochondria are stained only with some reagents and PFA fixation. Methanol fixation leads to the most consistent results: virtually no signal is present in untreated cells and H2O2 leads to a signal in the nucleus as expected. This agrees with the western blots, where endogenous MARylation is very low when untreated cells are lysed and analysed. It is possible that during cell lysis when mitochondrial contents are released, a highly active hydrolase reverses the modification of the mitochondrial proteins detected using IF. The low signal in western blot with lysates from unstimulated cells, argues that either the ARTs are present in inactive form and need a stimulus to activate them, or that hydrolases are highly active.

The observed lack of MARylation in the cytosol is unexpected as many of the mono-ARTs are expressed in the cytosol (11, 12, 20). The question remains which stimuli activate these enzymes in cells: is there a generic signal that can activate them all, analogous to the caspases being activated in a cascade during regulated cell death? Or are there specific triggers that activate specific ARTs, such as possibly viral infection activating the interferon-responsive ARTs, high levels of unfolded proteins activating PARP16, or translational shutdown activating the stress granule-associated ARTs, et cetera. The alternative explanation for the low MARylation signal is that the corresponding hydrolases are highly active and prevent visualisation of the MARylation due to high turnover rates. Increased knowledge about the hydrolases, as well as specific inhibitors or knockout cells are required to distinguish between these two possible explanations for lack of detectable endogenous MARylation. The availability of several reagents capable of detecting MARylation on *in vitro* modified proteins, in cell lysates as well as for microscopy, will enable future work studying these regulatory mechanisms. Likewise, some of the available reagents are suitable for the detection of MARylated RNA, which has thus far only been studied *in vitro*. It is not clear to which extent the *in vitro* assays represent the activity of the ARTs in cells: the sheer amounts of enzymes and substrates being brought together may induce artificial modification of exposed residues. Measuring in cell ADP-ribosylation will likely provide much more accurate information than *in vitro* assays.

One of the major outstanding questions in the field is thus to identify the signals which trigger endogenous ADP-ribosyltransferase activity in cells, as well as the extent wherein hydrolases reverse the modification in cells and during lysis. This work highlights the importance of careful sample preparation to avoid both loss and artificial gain of MARylation signal and provides a comparison of the currently available reagents, which will allow researchers to make an informed decision as to which reagent to use for their specific purposes.

## METHODS

### Plasmids

Many plasmids used to express proteins were gifts from other labs, as indicated below (**Table 1**). For expression of GFP-ARTs in mammalian cells, we amplified the different genes using appropriate primers for the full-length gene products from HeLa cDNA where possible or used gBlocks from IDT. These were either transferred into the Gateway system. All ARTs harbour the GFP-tag on the N-terminus in these constructs. Full sequences are available with the plasmids on Addgene. The generation of pDONR-mPARP14-m2m3 constructs was described before (43). These we transferred using the Gateway system to a Gateway-compatible pEGFP plasmid or amplified with appropriate primers for restriction cloning into pET28a.

**Table 1.** Plasmids used in this study.

| Plasmid name | Vector | Insert | Ref. | Addgene |
| --- | --- | --- | --- | --- |
| pNH-TrxT-MACROD1_77-325 | pNH-TrxT | MACROD1_77-325 | (58) | na |
| pDEST17-MACROD2 | pDEST17 | MACROD2 | (13) | 172593 |
| pDEST17-TARG1 | pDEST17 | TARG1 | (59) | 172594 |
| pNH-TrxT_PARGcat_448-976 | pNH-TrxT | PARGcat_448-976 | (60) | na |
| pDEST17-DXO | pDEST17 | DXO | This study | 177993 |
| pDEST17-DXO_E253A | pDEST17 | DXO_E253A | This study | 177994 |
| pNIC-Bsa4_ARH1 | pNIC-Bsa4 | ARH1 | (60) | na |
| pDEST17-ARH3 | pDEST17 | ARH3 | This study | 183051 |
| pGST-PARP10_818-1025 | pGST | PARP10_818-1025 | (6) | na |
| pASK60-mART2.2-FlagHis6x | pASK60 | Mouse ART2.2 | (61) | na |
| pET15b-rPtxS1 | pET15b | rPtxS1 | (49) | 173076 |
| pGEX-C3 bot | pGEX | Clostridium botulinum C3 toxin | (62) | na |
| pET28a-mPARP14-macro2/3-Fc | pET28a | Mouse PARP14-macro2/3 | This study |  |
| pET28a-mPARP14-macro2/3-GE-Fc | pET28a | Mouse PARP14-macro2/3-GE | This study |  |
| pEGFP-mPARP14-macro2/3-Fc | pEGFP-C1 | Mouse PARP14-macro2/3 | This study |  |
| pEGFP-mPARP14-macro2/3-GE-Fc | pEGFP-C1 | Mouse PARP14-macro2/3-GE | This study |  |
| pME_CD8L_FLAG-ART2.2GPI | pME | Mouse ART2.2 | (63) | na |
| pEGFP-PARP# | pEGFP | PARP1, PARP4, TNKS1, TNKS2, PARP8, PARP9, PARP13, PARP16 | (20) | na |
| pcDNA5-FRT/TO-N-mEGFP-PARP3 | pcDNA5-FRT/TO-N-mEGFP | PARP3 | This study | 178007 |
| pcDNA5-FRT/TO-N-mEGFP-PARP6 | pcDNA5-FRT/TO-N-mEGFP | PARP6 | This study | 178005 |
| pcDNA5-FRT/TO-N-mEGFP-PARP7 | pcDNA5-FRT/TO-N-mEGFP | PARP7 | This study | 178004 |
| pcDNA5-FRT/TO-N-mEGFP-PARP10 | pcDNA5-FRT/TO-N-mEGFP | PARP10 | This study | 177997 |
| pcDNA5-FRT/TO-N-mEGFP-PARP11 | pcDNA5-FRT/TO-N-mEGFP | PARP11 | This study | 177998 |
| pcDNA5-FRT/TO-N-mEGFP-PARP12 | pcDNA5-FRT/TO-N-mEGFP | PARP12 | This study | 177999 |
| pcDNA5-FRT/TO-N-mEGFP-PARP15 | pcDNA5-FRT/TO-N-mEGFP | PARP15 | This study | 178000 |

### Protein purification

The majority of ARTs used in this study were purified from bacterial expression systems with N-terminal His or GST-tags, with the exception of PARP1 which was produced as GFP-fusion protein from HEK293T cells as described below. As we found that many of the proteins are toxic to the bacteria, we optimised expression conditions and bacterial strains for each protein separately, with the optimal conditions summarised below **(Table 2**).

**Table 2:**
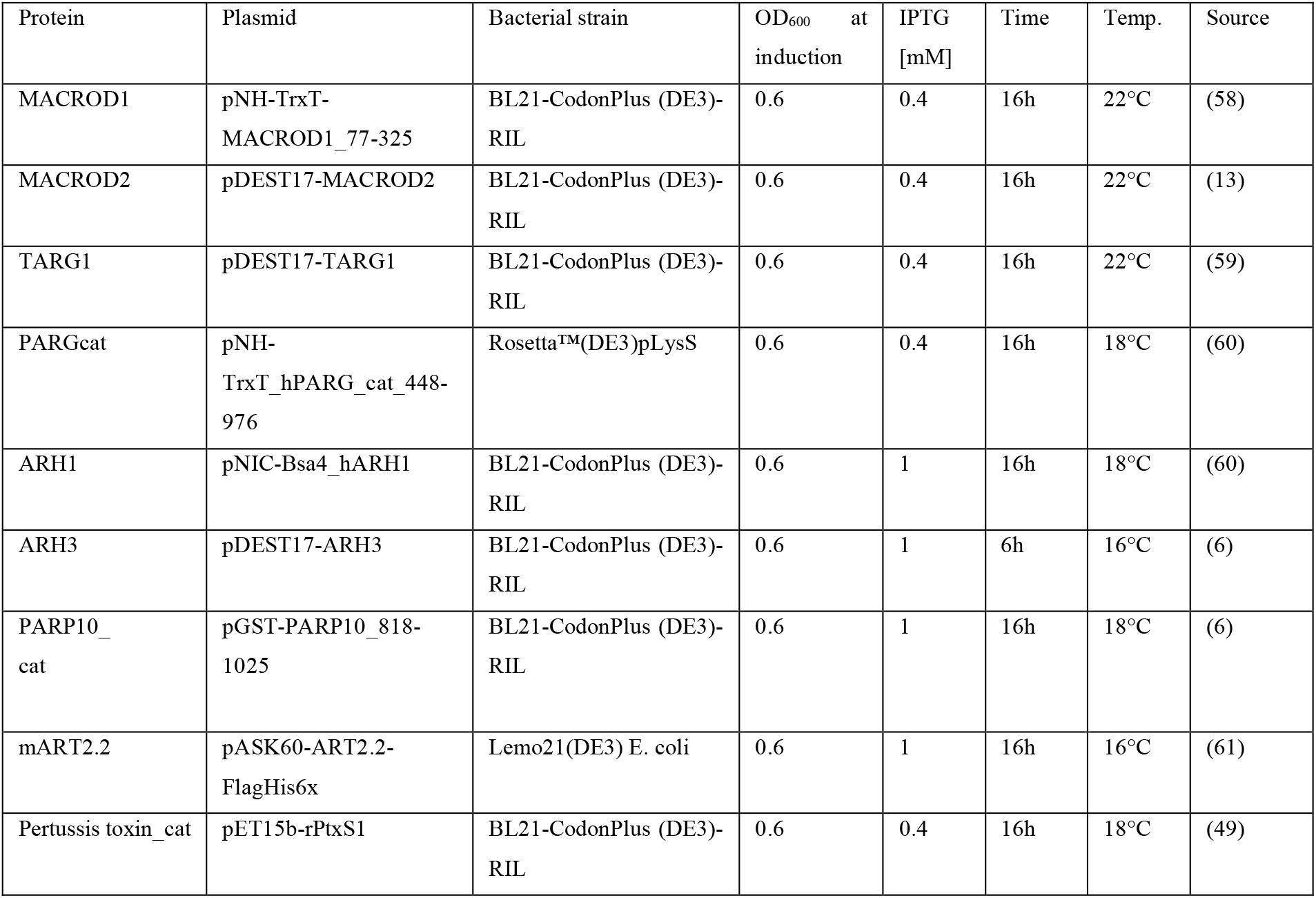
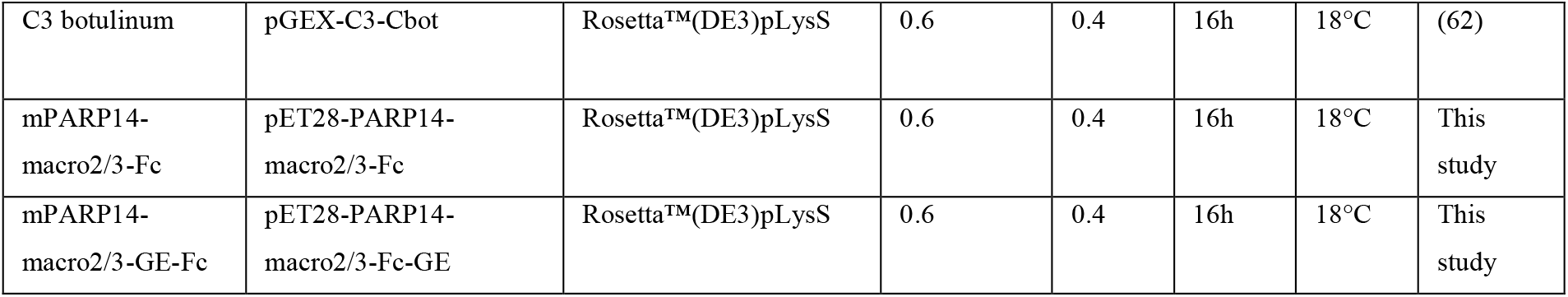
conditions for recombinant protein expression.

Bacterial cultures were spun down at 6,000 x *g* for 15 minutes at 4°C, followed by resuspension in lysis buffer. Cell suspensions were sonicated on ice using a Digital Sonifier 250 Cell Disruptor (Branson). Depending on the expressed protein construct and the pellet size, sonication varied in a range of 2 to 5 minutes at 15–20%, 30 seconds on, 40 seconds off. The cell lysates were centrifuged at 45,000 x *g* for 45 minutes at 4°C, to remove cell debris. The supernatant was used for affinity purification, using either glutathione sepharose 4G beads (GE Healthcare) or TALON metal affinity resin (Takara), for GST-tagged and His-tagged constructs, respectively, followed by dialysis. The composition of lysis, wash, elution and dialysis buffer was adjusted according to each proteins’ tag, properties and isoelectric point and is summarised in Table 3.

**Table 3.**
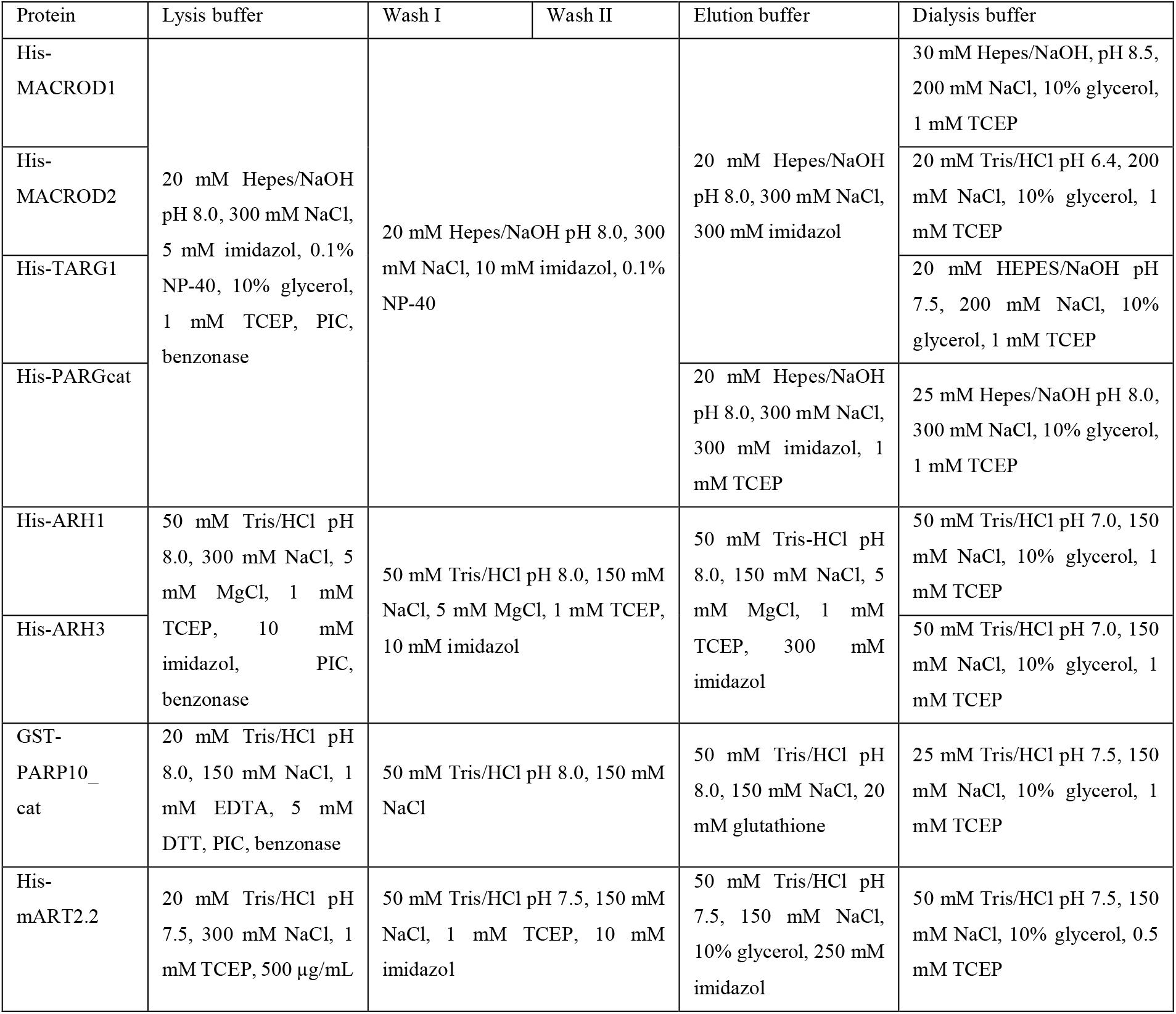
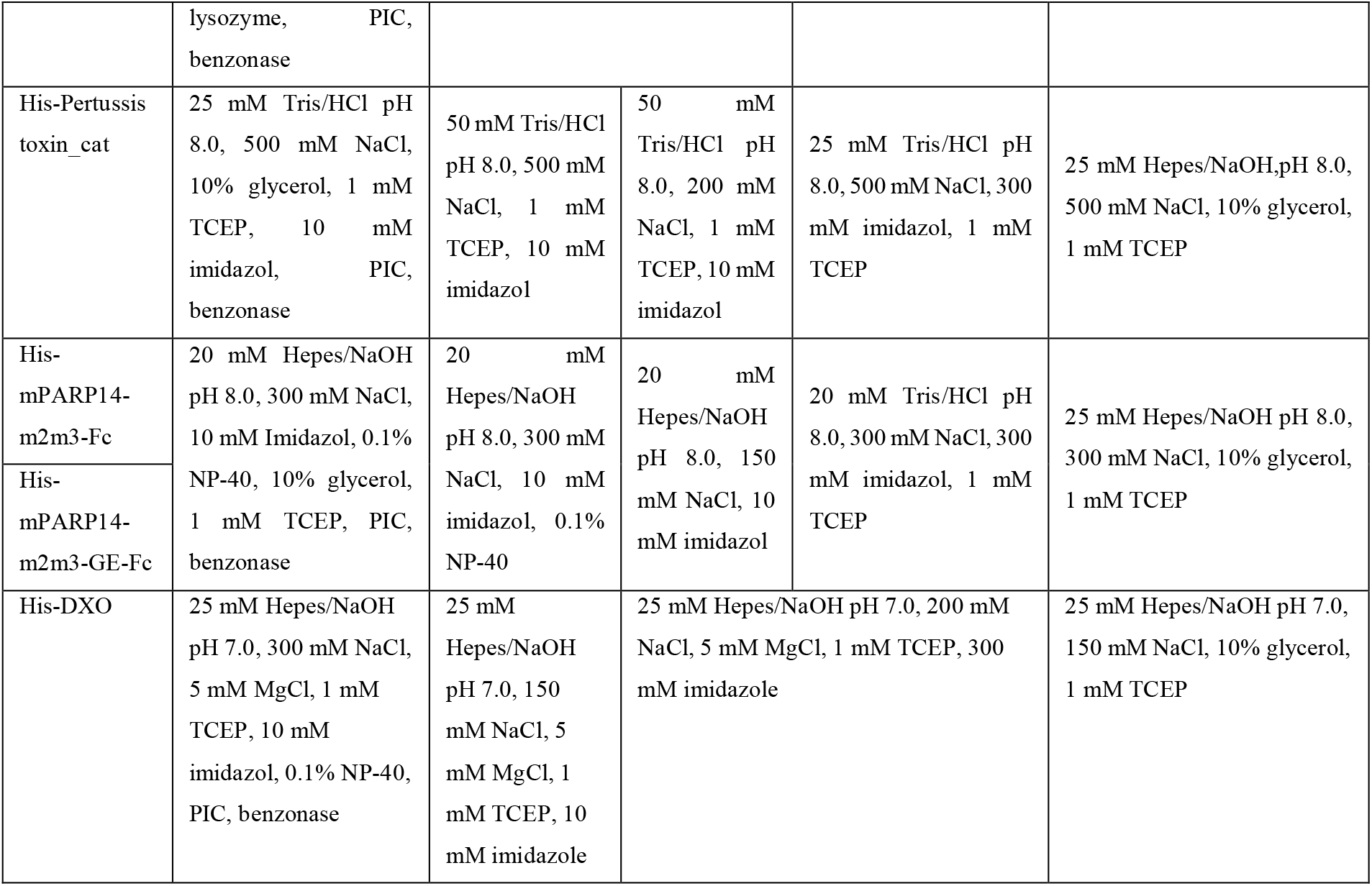

**Table 4.**
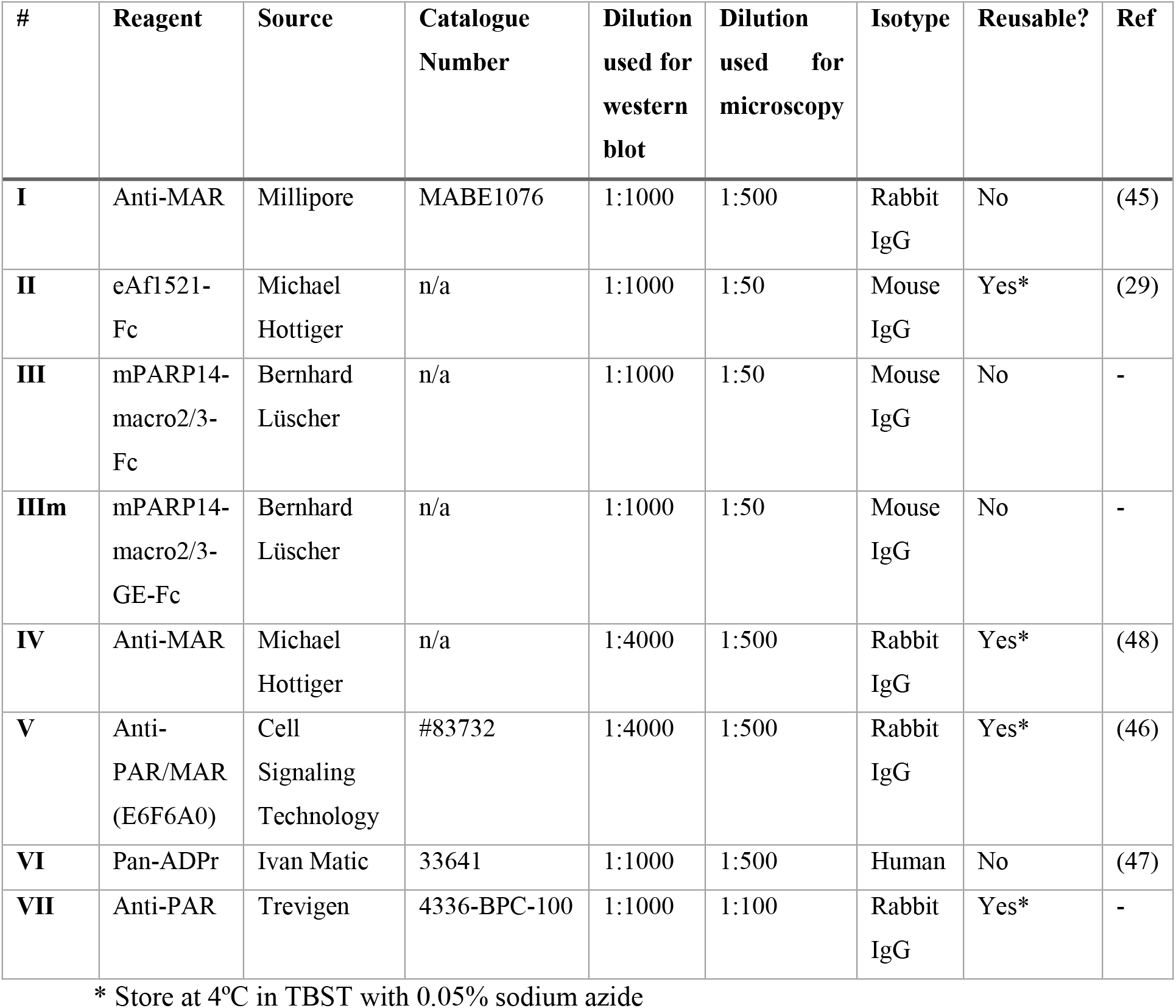
Antibody dilutions.

#### Mammalian cell culture and transfection

All cells were cultured in DMEM with pyruvate, 4.5g/L glucose (Gibco) and 10% heat-inactivated fetal calf serum (Gibco) in a humidified atmosphere with 5% CO2. Cells are routinely tested for mycoplasma contamination and confirmed negative at the moment of experiments. HEK293T cells were transfected using the calcium phosphate method as described in a step-by-step protocol available online (64). Briefly, cells were seeded in 6-well plates with 3 × 10^5 cells per well and transfected next day with 4 µg DNA per well. Approximately 6 hours after transfection, cells were washed and fresh full DMEM was added. 24 to 48 hours after transfection cells were lysed.

### Immunoprecipitation

GFP-PARP1 for enzymatic assays was immunoprecipitated from HEK293 cells. The cells were transfected with a plasmid encoding for GFP-PARP1 in 10 cm dishes and were further processed 24-48 hours after transfection. Cells were washed in warm DMEM without FCS and lysed in CoIP buffer (10 mM HEPES pH 7.5, 50 mM NaCl, 30 mM sodium Na4P2O7, 50 mM NaF, 0.2 % triton X-100 and 10% glycerol) containing protease inhibitor cocktail (8340, Sigma) and benzonase 1:10.000 (9025-65-4, Santa Cruz). Lysates were centrifuged at 4°C and maximum speed for 15 minutes to remove insoluble material, followed by incubation of the cleared supernatant with 10 µL prewashed GFP-coupled magnetic agarose (Chromotek) per 10 cm dish. After half an hour under rotation in the cold room, beads were washed in CoIP buffer, followed by a final wash in 1x PARP assay buffer. The material from 1 dish was split into 4 tubes for subsequent enzymatic reactions. Samples were processed immediately or stored in one-use aliquots at -20°C.

### Generation of cytosolic extracts

To generate substrate protein for some of the transferases and toxins, cytosolic extracts were generated. HEK293T cells were grown on 15 cm dishes until fully confluent and collected by trypsinisation. Cells were washed in hypotonic buffer (250 mM sucrose, 2 mM HEPES, 0.1 mM EGTA, pH 7.4), followed by incubation for approximately 15 minutes on ice to allow swelling, as monitored using a microscope. Cells were then broken by 20 strokes with a Dounce pestle B, followed by centrifugation at 11,000 x *g* to remove unbroken cells, nuclei and mitochondria. Resulting extracts are devoid of transferase PARP1, as well as devoid of the majority hydrolases, which are either mitochondrial, nuclear or expressed at low levels (65, 66).

### ADP-ribosylation and hydrolase assays

Protein ADP-ribosylation assays were routinely carried out at 37°C for 30 minutes unless indicated otherwise. Reactions were carried out in 30 µL containing 50 mM Tris-HCl, pH 7.5, 0.5 mM DTT, 0.1% Triton X-100, 5 mM MgCl2 and 50 µM β-NAD^+^ (Sigma) and 1 µCi [^32^P]-β-NAD^+^ (Hartman Analytics). 15 µl HEK293T cytosolic extract was used for the toxin and mART2.2 reactions. The amounts of enzymes used varied depending on the activity of the enzyme studied. Reactions were placed on ice and where possible, transferase inhibitors were added before adding hydrolases. For PARP10, OUL35 was used at 2 µM, for PARP1 olaparib was added at 5 µM. For mART2.2 and the toxins, no inhibitors were available and hydrolases were added to the cooled reaction. Hydrolase reactions were incubated at 37°C for 30 minutes, stopped by adding 4x SDS sample buffer, heated for 10 minutes at 60°C and run on SDS-PAGE. If protein coupled to a GFP-trap was analysed, we incubated the beads for 5 minutes at 95°C before loading on gel. Gels were dried and incorporated radioactivity was analysed by exposure of the dried gel to X-ray film.

For RNA modification, 7 µM of ssRNA were incubated with 2 µM TRPT1 in ADPr-buffer (20 mM HEPES-KOH, pH 7.6, 50 mM KCl, 5 mM MgCl2, 1 mM DTT, 500 µM NAD, 40U RNase inhibitor) at 37°C for 60 minutes while shaking. TRPT1 was digested by Proteinase K treatment (20 U), for 20 minutes at RT, before purification using the Monarch® RNA Cleanup Kit (T2030). Concentration was determined by spectrophotometric measurements (NanoDrop ND-1000 Spectrophotometer).

### Antibodies, western blot and slot blot

Cells were washed in warm DMEM without FCS before lysis to avoid unnecessary stress. Cell protein extractions were performed using RIPA buffer (150 mM NaCl, 1% Triton X-100, 0.5% sodium deoxycholate, 0.1% SDS, 50 mM Tris-HCl (pH 8.0)) supplemented with protease inhibitor cocktail, benzonase and olaparib. Proteins were separated on 10–15% gels and blotted onto nitrocellulose membranes using a BioRad TurboBlot apparatus using the high-molecular weight 10-minute program. A step-by-step protocol for our western blotting procedure is available online (67). Membranes were blocked with 5% non-fat milk in TBST for 30–60 minutes at RT, primary antibodies were diluted in TBST as indicated below and incubated overnight at 4°C, secondary antibodies were diluted 1:5000 in 5% non-fat milk in TBST and incubated for 30–60 minutes at RT. Some of the antibodies could be stored and reused multiple times as indicated in Table 2. Multiple wash steps were performed in between and after antibody incubations with TBST at RT for at least 5 minutes. Chemiluminescent signals were detected by either exposure to film or using the Azure600. Additional antibodies used: anti-GFP 1:2500 (600-101-215 Rockland), anti-HSP60 1:2500 (12165 Cell Signaling Technology), anti-PARP1 1:1000 (1 835 238 Roche), anti-tubulin 1:5000 (B-5-1-2 Santa Cruz). For slot blotting, 2µM of each peptide was slot blotted using a PR648 Slot Blot Blotting Manifold (Hoefer) onto nitrocellulose membrane. Subsequently, the membrane was processed identical to the described processing of western blots and detected using exposure to film.

### RNA and peptide slot blot

RNA was blotted on Hybond-N membrane (Amersham) using a PR648 Slot Blot Blotting Manifold (Hoefer). The membrane was activated in SSC-buffer pH 7.0 (150 mM NaCl, 15 mM NaCit) for 5 minutes. After assembling of the blotting sandwich, slots were flushed with SSC-buffer. 40 ng RNA in 200 µL SSC-buffer per slot was applied to the membrane. The membrane was air dried and samples were cross-linked using UV light (120 milijoules/cm2). The membrane was blocked with 5% non-fat milk in PBST for 60 minutes at RT, primary antibodies were diluted in PBST as indicated and incubated overnight at 4°C, secondary antibodies were diluted 1:5000 in 2% non-fat milk in PBST and incubated for 30 minutes at RT. Multiple wash steps with PBST were performed after both antibody incubations for at least 5 minutes. Chemiluminescent signals were detected using the Azure600. Peptides were blotted onto nitrocellulose activated in water, followed by blocking in 5% non-fat milk in PBST and antibody incubations. Antibody dilutions and wash steps were identical to the processing of western blots.

### Confocal microscopy

U2OS or HeLa cells were seeded onto glass coverslips in 24 well plates. For H2O2 treatment, medium was removed and replaced with warm PBS containing 1mM H2O2 for 5 minutes. Cells were washed once in warm DMEM without FCS and fixed for 20 minutes with 4% paraformaldehyde in PBS at room temperature. Alternatively, ice-cold methanol was added to the cells on coverslips after removal of the growth medium, followed by 5 minutes incubation on ice and subsequent quick washing in PBS. For glyoxal fixation glyoxal solution was added to the cells for 30 minutes on ice followed by 30 minutes at room temperature. After washing in PBS, the samples were quenched using 100 mM ammonium chloride and finally permeabilization using 0.1% Triton X-100. Regardless of fixation method, subsequent blocking was done in PBS supplemented with 1% BSA in PBS with 0.1% Triton X-100 for 1 hour at RT. Primary and secondary antibodies were applied for 1 hour, with extensive washing in between. Primary antibody dilutions are indicated (**Table 4**). Secondary antibodies used: AlexaFluor594 anti-mouse, AlexaFluor594 anti-rabbit, AlexaFluor633 anti-human (all ThermoFisher) used 1:2000 in PBS with 0.2% BSA. Following extensive washing in PBS and demineralised water, coverslips were mounted on microscopy slides using Prolong Anti-Fade Diamond Mountant containing DAPI (ThermoFisher). The samples were analysed with a Zeiss LSM710 Confocal Laser Scanning Microscope equipped with an AxioCam (Zeiss) and a C-Apochromat 20x objective. Step-by-step immunofluorescence protocols are available online (68).

## DATA AVAILABILITY STATEMENT

Unprocessed blots are presented either in the manuscript or in the Supplementary Data Files. Expression constructs generated in this study will be submitted to Addgene. We are also able to ship the recombinant proteins we generated to labs interested.

## AUTHOR CONTRIBUTIONS

L.W., G.A., T.B. and J.M. purified proteins; L.W. and R.Ž. performed RNA slot blots; T.B., J.M. and K.F. performed radioactivity experiments; K.F. performed western blots; K.F. and R. Ž. Performed confocal microscopy; G.A. and R.Ž. cloned expression constructs; M.B., A.G. and B.L. generated the murine Parp14-m2m3-Fc fusion constructs; K.F. and R. Ž. conceived this study, designed the experiments and analysed the data; R.Ž., B.L. and K.F. acquired funding; K.F. wrote the manuscript with input from R.Ž. and L.W. All authors have proof-read and agreed with the submission of the manuscript.

## ACKNOWLEDGEMENTS

We thank the researchers in the ADP-ribosylation community who have put their expression constructs and detection reagents at our disposal to allow thorough characterisation of these materials: Michael O. Hottiger provided the eAf1521 and ADPr antibody; Ivan Matic provided a pan-ADPr antibody. The SpvB, mART2.2 bacterial and human expression plasmids were a gift from Fritz Koch-Nolte; truncated pertussis toxin, PARG and ARH1 constructs were kindly provided by Lari Lehtiö; Paul Chang provided GFP-PARP expression constructs. Purified recombinant HPF1 was provided by Patricia Korn. We are grateful for helpful discussions with and suggestions from Michael O. Hottiger and Michael Cohen. This work was supported by the Confocal Microscopy Facility, a Core Facility of the Interdisciplinary Center for Clinical Research (IZKF) Aachen within the Faculty of Medicine at RWTH Aachen University. Funding was provided by the START program of the Medical Faculty of RWTH Aachen University to K.F. (10/18) and R.Ž. (13/20) and by the German Research Foundation DFG to B.L. (LU466/16-2) and K. F. (FE1423/3-1).

## Notes

### Competing Interest Statement

The authors have declared no competing interest.

