## Supplementary Data for "Systematic analysis of ADP-ribose detection reagents and optimisation of sample preparation to detect ADP-ribosylation *in vitro* and in cells"

**Supplementary Figure 1: generation of specific substrates**

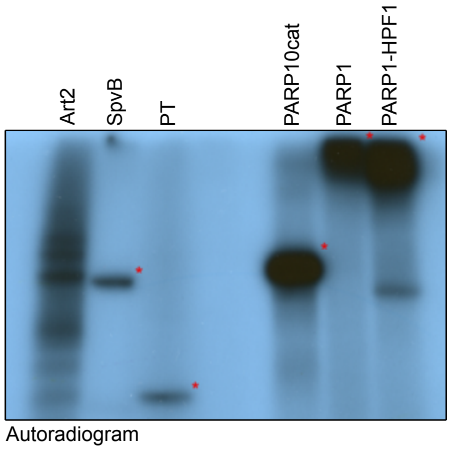


**Supplementary figure 1**: Indicated ADP-ribosyltransferases were incubated with $^{32}$P-labeled NAD$^+$ for 30 minutes and analysed using SDS-PAGE and exposure to film. Art2 and SpvB modify substrates in a cytosolic extract whereas the truncated pertussis toxin (PT), PARP10 catalytic domain, PARP1 and PARP1/HPF1 automodify. Identical amounts of this reaction were loaded on SDS page and blotted to compare the reagent specificity on identical substrates.

**Supplementary Figure 2: Coomassie and film belonging to figure 1**

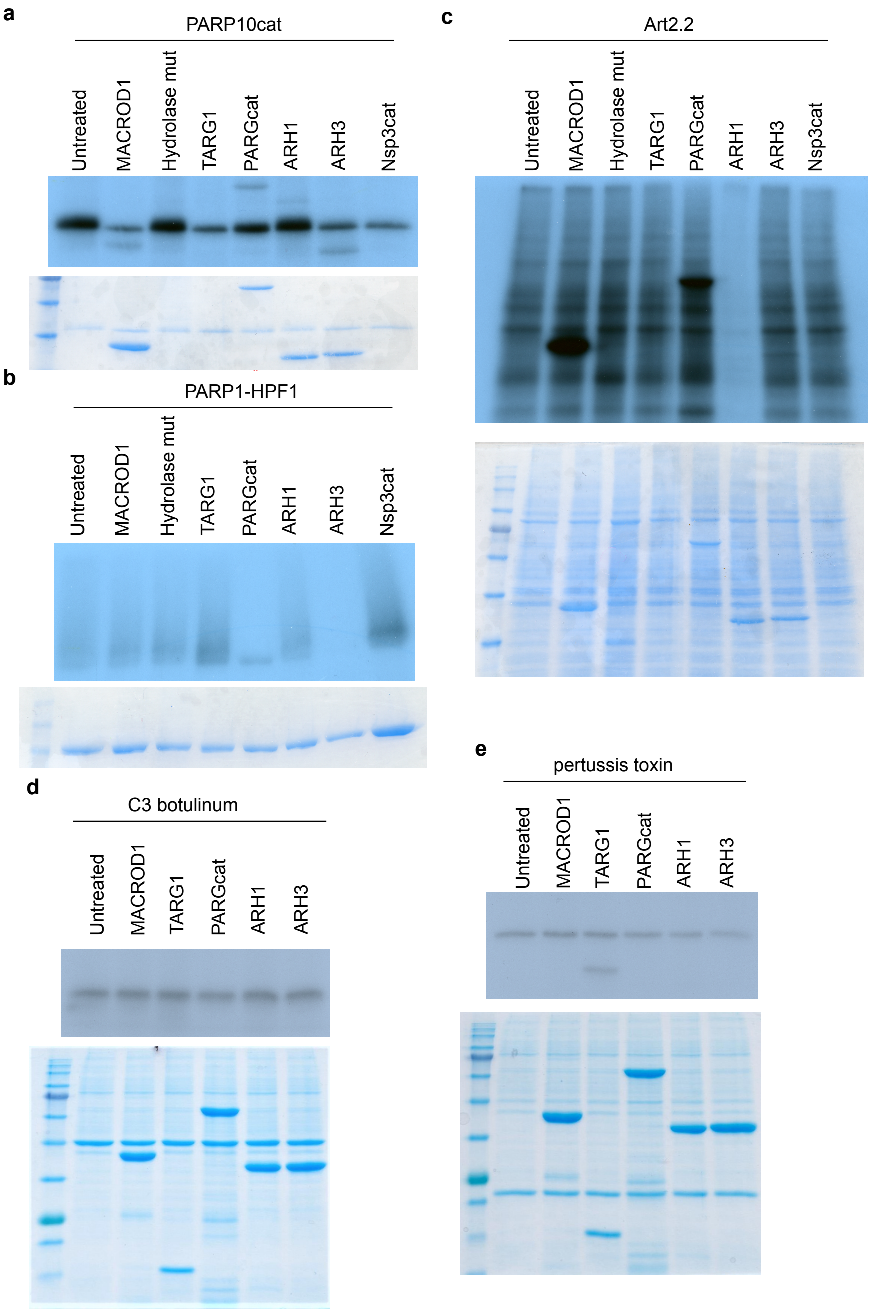

**Supplementary Figure 3: PARP1/HPF1 ADP-ribosylation is reduced by PARG but not MACROD1**

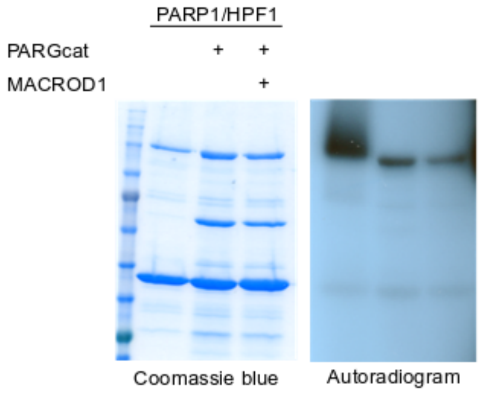


**Supplementary figure 3**: Immunoprecipitated GFP-PARP1 was supplemented with recombinant His-HPF1 and automodified using $^{32}$P-NAD$^{+}$ in an ADP-ribosylation reaction for 30 minutes. The automodified PARP1 was further incubated alone, with PARGcat or PARGcat and MACROD1. The signal present after PARG and MACROD1 treatment should represent MARylation on serines, which is not reversed by MACROD1.

**Supplementary Figure 4: some antibodies lead to high background signal on slot-blotted peptides**

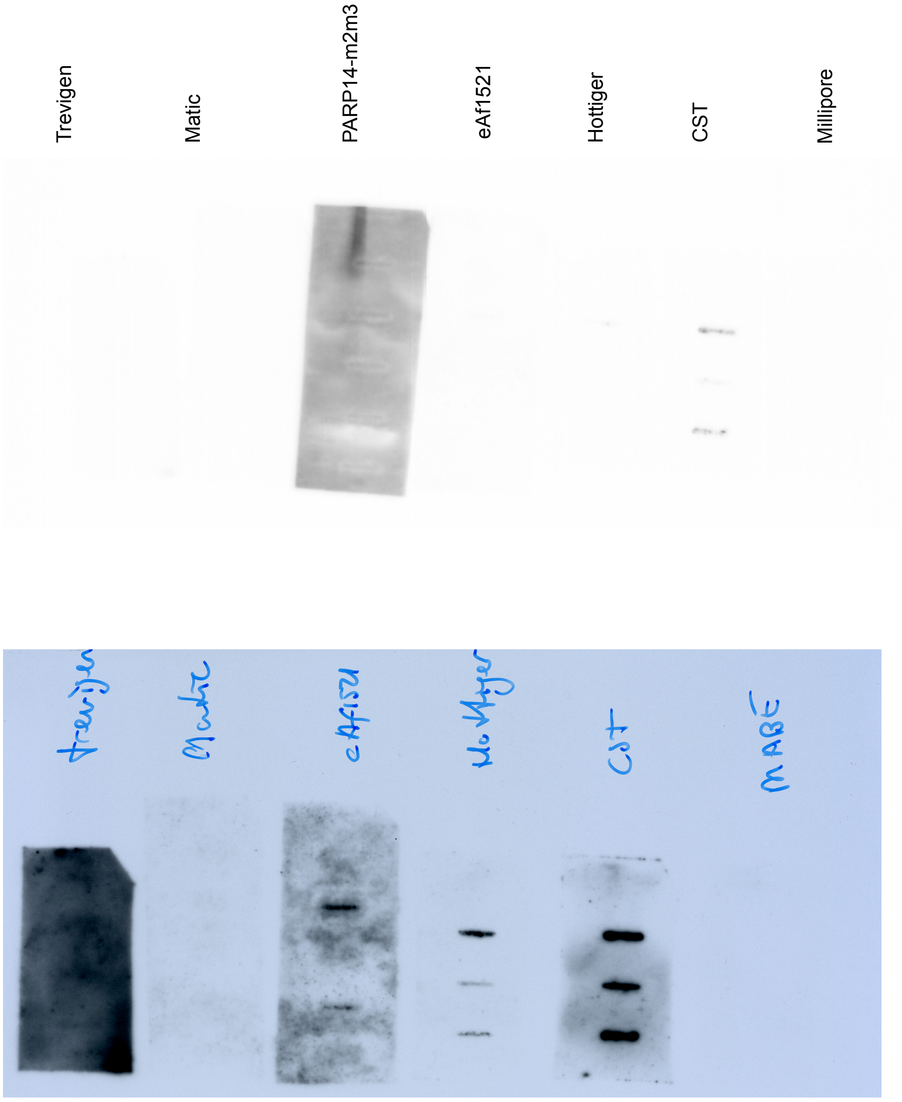


**Supplementary Figure 4:** slot blots with different chemically modified peptides were analysed using the indicated reagents. Blots were first detected using the Azure system (top). The PARP14-m2m3 strip was removed due to high background, before subsequent film detection of the strips (bottom).

**Supplementary Figure 5: Quantification and different exposures of the thermolability of ADPr-peptides**

**a**

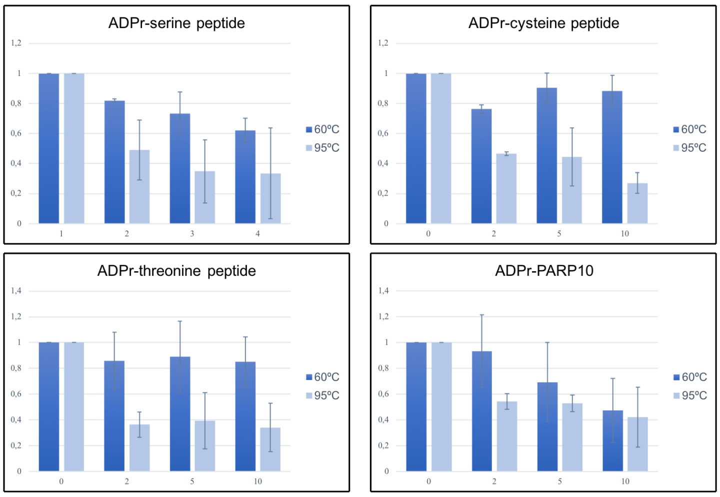


**b**

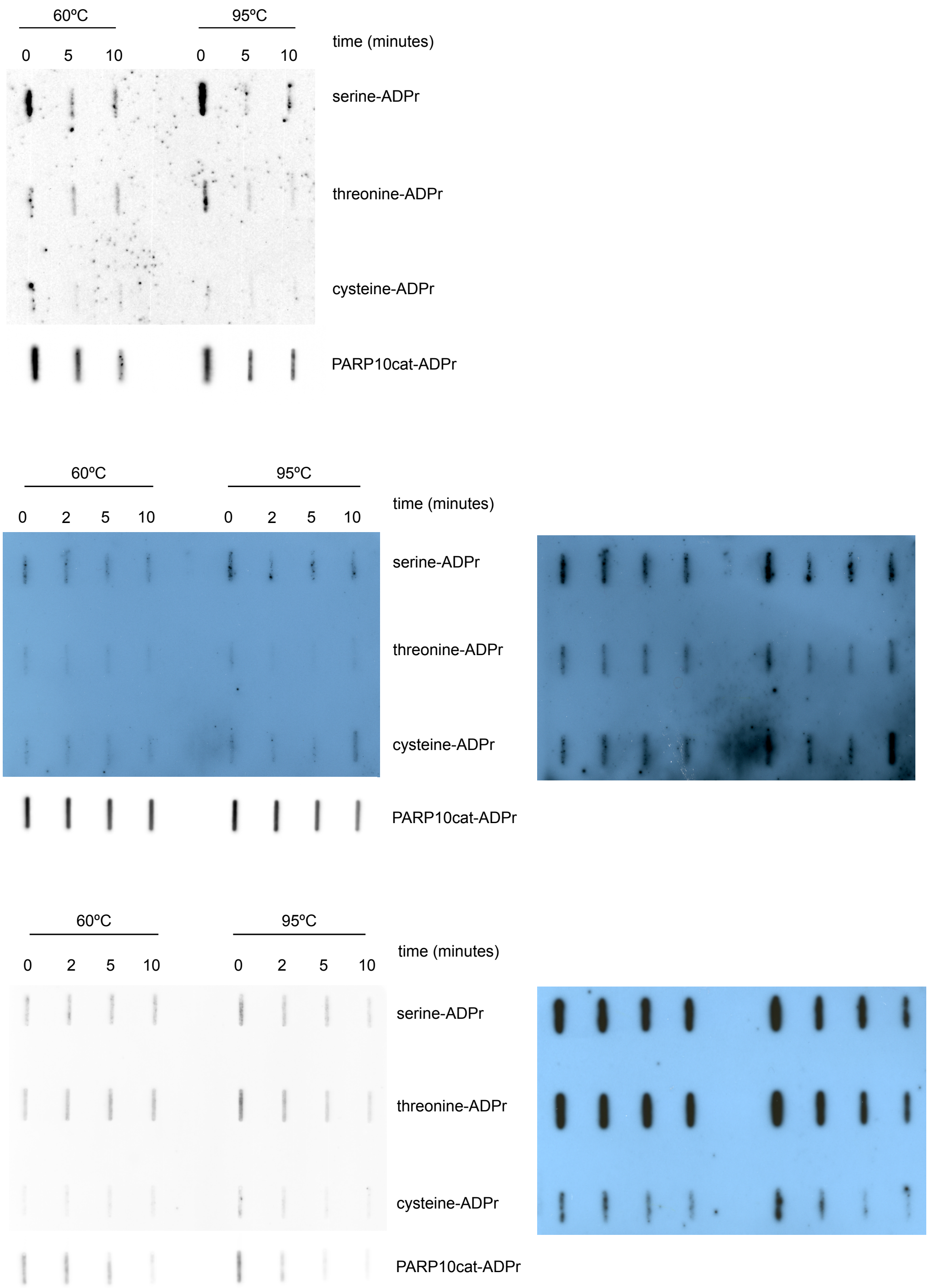


**Supplementary figure 5:** different exposures of the technical replicates of slot-blots with peptides incubated at different temperatures. Exposures were chosen were the strongest signal is not oversaturated for quantification. Images were converted to 8-bit and quantified using the magic wand tool in Fiji. The PARP10 signals were much stronger than the peptides and detected using Azure equipment, the weaker signal from the peptides was in addition detected using film.

**Supplementary Figure 6: Some reagents detect adenylylated RNA and reagent IV detects AMPylated proteins in addition to ADP-ribosylated proteins**

**a**

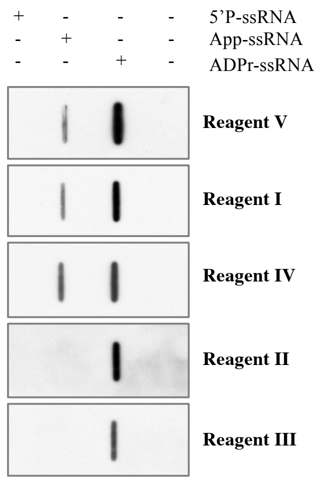


**Supplementary Figure 6. (a)** ssRNA was adenylylated using a commercial kit and subsequently slot-blotted. ADPr-ssRNA was used as positive control. Blots were analysed using indicated reagents.

**Supplementary Figure 7: immunofluorescence staining using the different reagents**

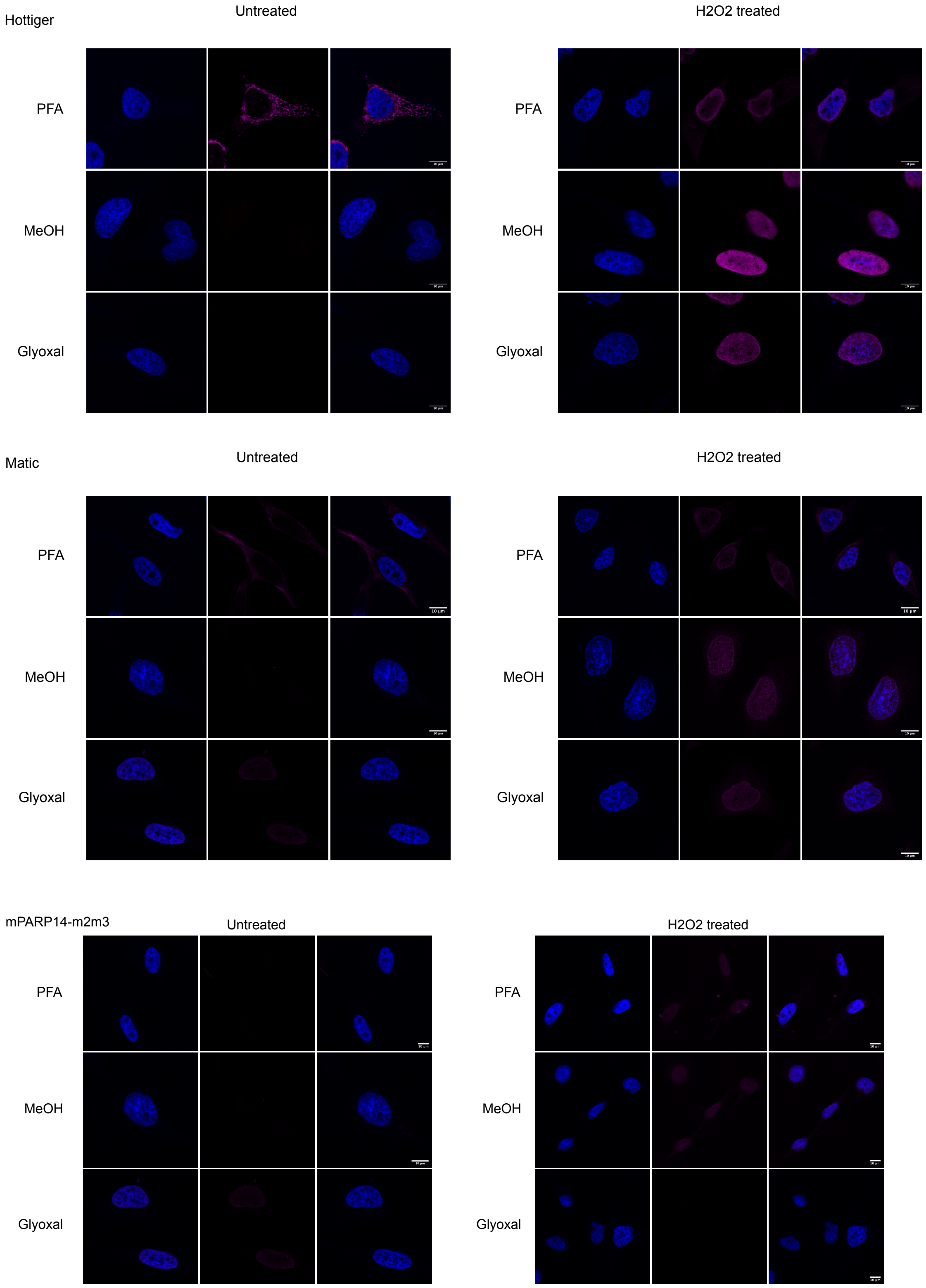

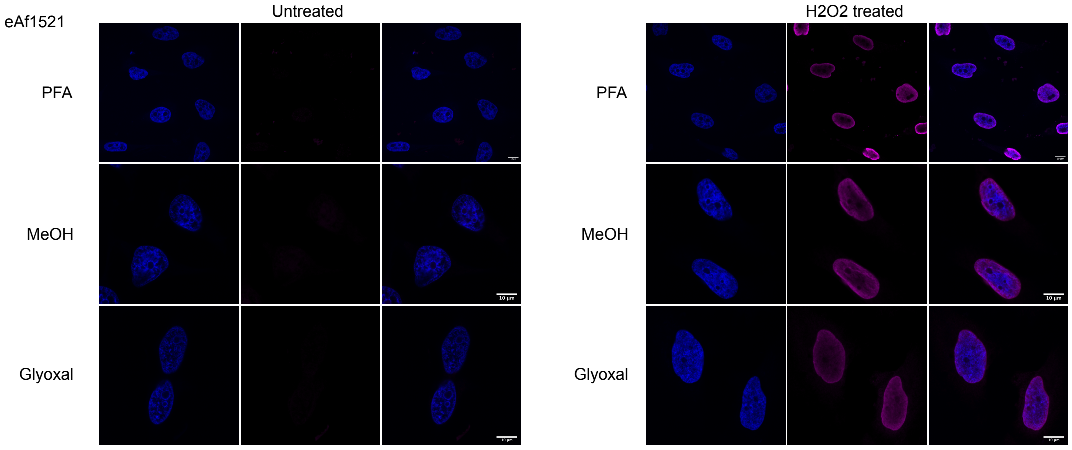


**Supplementary figure 7:** HeLa cells were grown on coverslips and fixed using either 3.7% paraformaldehyde, ice-cold methanol or glyoxal, followed by staining using the indicated antibodies and DAPI. Cells were imaged using confocal microscopy.

**Supplementary Figure 8 unprocessed blots with size marker overlay**

Unprocessed blot belonging to figure 3

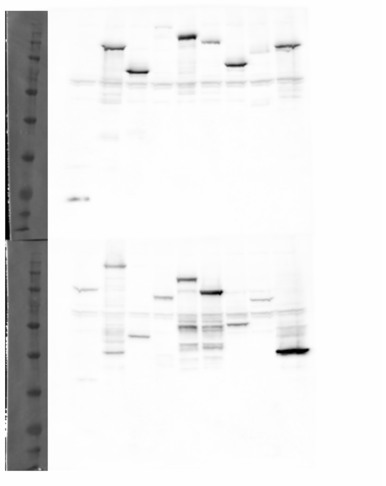

Unprocessed blots belonging to figure 4

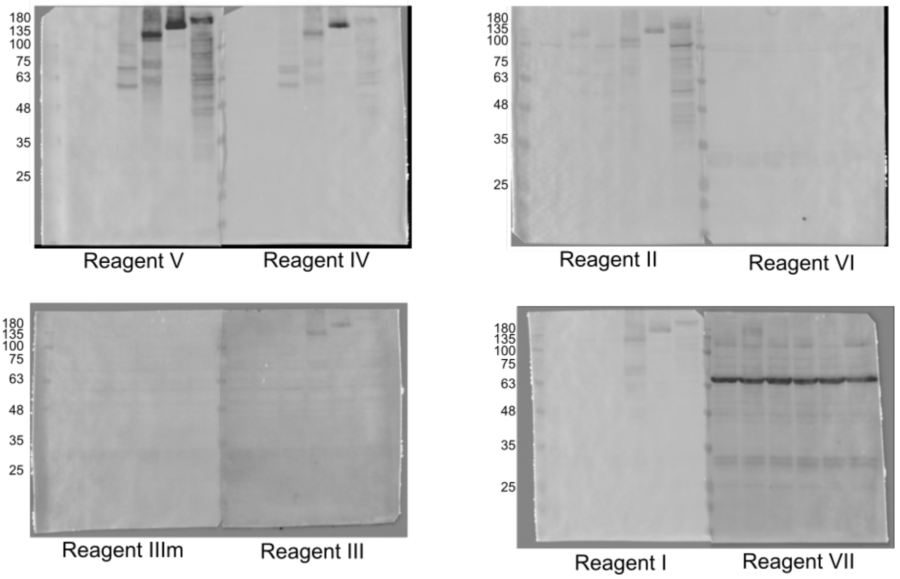
